# Differential genetic expression within reward-specific ensembles in mice

**DOI:** 10.1101/2023.11.02.565378

**Authors:** Carl G. Litif, Levi T. Flom, Kathryn L. Sandum, Skylar L. Hodgins, Lucio Vaccaro, Jerry A. Stitzel, Nicolas A. Blouin, Maria Constanza Mannino, Jason P. Gigley, Todd A. Schoborg, Ana-Clara Bobadilla

## Abstract

Maladaptive reward seeking is a hallmark of cocaine use disorder. To develop therapeutic targets, it is critical to understand the neurobiological changes specific to cocaine-seeking without altering the seeking of natural rewards, e.g., sucrose. The prefrontal cortex (PFC) and the nucleus accumbens core (NAcore) are known regions associated with cocaine- and sucrose-seeking ensembles, i.e., a sparse population of co-activated neurons. Within ensembles, transcriptomic alterations in the PFC and NAcore underlie the learning and persistence of cocaine- and sucrose-seeking behavior. However, transcriptomes exclusively driving cocaine seeking independent from sucrose seeking have not yet been defined using a within-subject approach. Using Ai14:cFos-TRAP2 transgenic mice in a dual cocaine and sucrose self-administration model, we fluorescently sorted (FACS) and characterized (RNAseq) the transcriptomes defining cocaine- and sucrose-seeking ensembles. We found reward- and region-specific transcriptomic changes that will help develop clinically relevant genetic approaches to decrease cocaine-seeking behavior without altering non-drug reward-based positive reinforcement.

## Introduction

Cocaine use disorder (CUD) is characterized by the recurrent seeking of cocaine after periods of abstinence and withdrawal, often leading to relapse (*1*). The absence of pharmaceutical intervention for attenuating cocaine seeking is partially due to a lack of understanding of the discrete molecular alterations driving the rewarding effects of cocaine apart from natural rewards, e.g., sucrose. Cocaine and sucrose seeking involve cross-talking reward-signaling regions: the nucleus accumbens core (NAcore) and prefrontal cortex (PFC) (*2–5*). Within the NAcore and PFC, sparse activity-dependent neural circuits, defined as neuronal ensembles (*6*), are linked to cue-associated cocaine and sucrose seeking and are mostly separate despite having an overlapping population of neurons (*7–9*).

Studies focused on ensemble-specific genetic expression in cocaine seeking often lack within-subject comparisons between different rewards. Mounting evidence indicates that contingent psychoactive intake facilitates persisting alterations in gene expression within the NAcore and PFC (*10–16*). However, little is known about gene expression profiles specific to ensembles activated during cocaine seeking after self-administration and extinction training.

Considering these gaps in our understanding of CUD, we focused on two main questions: Do neuronal ensembles linked with cocaine and sucrose seeking express different genes, or do they regulate the expression of the same genes differently? Do cocaine and sucrose seeking have similar or different gene expression in two critical regions of the reward system, e.g., the NAcore and PFC? To address these questions, we used a within-subject approach to characterize sex-considerate and region-dependent gene expression profiles for cocaine and sucrose seeking mice.

Using targeted recombination in active populations (Ai14:cFos-TRAP2) (*17*) and fluorescent activated cell sorting (FACS) (*15, 18*), we identified and isolated ensembles in the NAcore and PFC from temporally distinct cocaine- and sucrose-seeking events. We then completed unbiased bulk RNA-sequencing (RNAseq) to compare significant (false discovery rate (FDR) < 0.25) (*19, 20*) region-dependent and reward-specific genetic profiles for cocaine- and sucrose-seeking ensembles in reference to naïve mice. Using this approach, we developed a repository of sex-considerate genetic information to define distinct or intersecting genetic alterations for cocaine- and sucrose-seeking ensembles in the NAcore and PFC.

## Materials and Methods

### Ai14:cFos-TRAP2 Mice

Female and male Ai14:cFos-TRAP2 transgenic mice were obtained by crossing male tamoxifen-inducible cFos-Cre-recombinase knock-in mice (Fostm2.1(icre/ERT2)Luo/J, Strain# 030323; The Jackson Laboratory) with female Ai14 loxP-flank regulated Cre-reporter knock-in mice (B6;Cg-Gt(ROSA)26Sortm14(CAG-tdTomato)Hze/J; Strain# 007914; The Jackson Laboratory). Mice were individually housed on a 12:12 reverse light schedule during the length of dual-reward self-administration conditioning. Sample size and attrition rate were determined from dual-reward self-administration conditioning previously established methodology (*8*). All procedures were conducted in accordance with the IACUC and NIH animal handling procedures.

### Dual Reward Self-Administration, Extinction, And Reinstatement Tests

Both sexes were used for dual-reward self-administration of intravenous cocaine or oral sucrose intake. During self-administration acquisition, all mice were on a food-restricted diet (80% of normal food intake = 3.2 g) to increase reward-specific operant conditioning responses. Female and male 8-16-week-old mice (20-30g) underwent catheter implantation following a previously established methodology (*8*). All mice underwent dual-reward self-administration of intravenous cocaine infusions and oral intake of sucrose pellets during 2-hour daily sessions (10 days of cocaine reward and ten days of sucrose reward: 20 alternating days in total) within standard mouse modular test chambers. Each chamber included two nose-poke (NP) holes associated with cocaine infusions (0.5 mg/kg/inf; NIDA) or delivery of sucrose pellets (15 mg; Bio-Serv). Each NP was paired with an availability light cue (always on until mice associate with NP; off for ten seconds after NP activation) and an internal NP light cue (on for 3 seconds after NP activation). The reward-associated NP not used during a reward-specific session served as an inactive NP negative control (i.e., cocaine-associated NP during a sucrose session acted as an inactive NP with no cues or reward administration and vice versa). Following reward-specific cue conditioning, mice were extinguished of rewards or cues until reaching a 70% reduction of NP association compared to the averaged last three days of reward self-administration. Following the extinction phase, 30-minute reinstatement sessions for each reward were carried out with the correct reward-associated light cues and without administering the reward. Following intraperitoneal injection of 4-hydroxy-tamoxifen (4-OHT, 50 mg/kg, Sigma) directly after the cocaine reinstatement session, the Cre-regulated TRAP2 mechanism tagged active cocaine-seeking cells with endogenous tdTomato fluorescence. Following sucrose reinstatement sessions, mice were euthanized 60 minutes post-session to capture peak cFos expression reflective of active cells during sucrose-seeking behavior.

### Extraction of Reward-Specific Ensemble RNA

Whole brains were extracted from euthanized mice (5% isoflurane exposure followed by cervical dislocation) and incubated for 5 minutes on ice in cellular stabilizing solution: 45% HEBG medium (2% 50X B27 (Thermo Fisher; 17504044) and 0.25% 100X Glutamax (Thermo Fisher; 35050061) per 1 mL of Hibernate E-Minus Phenol Red (Thermo Fisher; NC1506837)), 45% artificial cerebral spinal fluid (aCSF; 7.13 g/L NaCl (Sigma-Aldrich; S9888-500G), 0.23 g/L KCl (Sigma-Aldrich; P3911-25G), 0.14 g/L MgSO4 (Sigma-Aldrich; 746452-500G), 0.19 g/L CaCl2 (Sigma-Aldrich; C33067-100G), 0.048 g/L NaH2PO4 (Millipore-Sigma; 10049-21-5), 2.1 g/L NaHCO3 (Millipore-Sigma; 144-55-8), 1 L diH2O adjusted for dry reagent volumes), 9.97% RNAlater (Thermo Fisher; AM7020), and 0.03% RNaseOUT (Thermo Fisher; 10777019). Brain sections (1.94-0.64 bregma; 1.5 mm thickness) containing the nucleus accumbens core and prefrontal cortex were cut using a brain matrix (0.5 mm spaced slits) kept cold at -20°C with ice-cold razors. Punches of brain tissue were made using a 1 mm biopsy punch for each nucleus accumbens bilateral region and two 1 mm sized punches for the prefrontal cortex. Brain regions from three mice per sex were pooled together for adequate RNA collection. Brain punches were immediately placed in enzymatic dissociation solution (2mg/mL papain (Worthington; LS003120) diluted in 1 mL HEBG with 0.125% 100x Glutamax – heated in 30C water bath for 10 minutes before immediately placing on ice for brain punches) on ice for 20 minutes. Mechanical trituration while incubating in papain solution was carried out using 10x syringe fluctuations with a 23G needle followed by a 27G needle. Caution is taken to not introduce air bubbles during syringe fluctuations that otherwise rupture cell membranes. Tissue was centrifuged for 3 minutes at 1000 rcf at 4°C, and the supernatant was carefully discarded via pipette, leaving ∼20 uL medium to prevent significant cell loss. Cells were slowly resuspended via pipette with 500 uL of 4°C HEBG medium and then filtered through a 40 um cell strainer. Semi-permeabilization was carried out by adding 500 uL of -20°C ethyl acetate (95% ethanol; 5% acetate) to the 500 uL of HEBG medium with cells for a final 50% ethyl acetate concentration. Tubes were slowly inverted by hand for 1 minute briefly before centrifuging for 5 min at 1000 rcf in 4°C. The supernatant was discarded carefully via pipette leaving ∼20 uL to prevent cellular loss and cells are resuspended gently via pipette in 500 uL of 4°C HEBG medium with neuronal (NeuN-Alexafluor-405; conc. 5:500; Novus Biologicals; NBP1-92693AF405) and cFos-labeling conjugated antibodies (cFos-Alexafluor-488; conc. 5:500; Novus Biologicals; NBP2-50037AF488). Cells were incubated at 4°C while rotating in the dark for 45 minutes. Cells were centrifuged for 5 minutes at 1000 rcf at 4°C, and the supernatant was carefully discarded via pipette, leaving ∼20 uL medium to prevent significant cell loss. Cells were resuspended in 1 mL of 4°C HEBG medium and filtered with a 40 um cell strainer into a sterile 5 mL FACS tube and kept at 4°C. Cells were then sorted for neuronal and reward-specific tags (neuron – immunolabeled NeuN; cocaine – endogenous tdTomato; sucrose – immunolabeled cFos; overlap – tdTomato/cFos) using FACS (FACSMelody; FACSChorus Software 100-um nozzle; BD Biosciences). 80k-100k events per reward-specific population were sorted directly into the RLT buffer (RNeasy Mini Kit; Qiagen; 74104) for downstream RNA extraction using RNeasy Mini Kit. RNA samples were kept at -80°C until downstream RNA sequencing.

### RNA Sequencing of Reward-Specific Neurons

RNA samples (average 260/280 ratio of 2.12) underwent library preparation and next-generation RNA sequencing at the University of Colorado Anschutz Medical Campus Genomics and Microarray Core. Low-input ribosome depleted (SMART-seq; Takara Bio; library size: 200-500bp) library prep was used for downstream paired-end 2x150bp RNA sequencing at a 50 million read depth using NovaSEQ 6000 (Illumina). Raw fastq output files were used for downstream bioinformatic analyses. Raw fastq and transcript count files are available on gene expression omnibus (GEO) under accession number GSE247029.

### Bioinformatic Analysis of Reward-Specific Transcriptomes

Raw sample files in fastq form were trimmed for Takara library prep adaptations using the “CutAdapt” (*21*) Linux-based program. Samples were mapped to mouse transcriptome (grc39) using Linux-based “Salmon” (*22*) program. Output counts from mapping were further normalized, linearly fit, and filtered to remove highly variable low expressed counts with “tximport” (*23*), “edgeR” (*24*), and “Limma” (*25*) R packages. DEG lists are represented in log2(fold change) format. “Glimma” (*26*) R package produced classical metric multidimensional scaling plots to compare sample variation. “Venndetail” (*27*) and “ggven” (*28*) R packages were used to compare reward- and region-specific DEGs that were significant (FDR < 0.25) with log2FC less than -0.75 or greater than 0.75. Heatmap representations of log2FC gene expression values were created using Prism 10 (GraphPad) and ordered sequentially based on the relevant significant specific DEG list (represented by *). Volcano plots were created using the “EnhancedVolcano” (*29*) R package to compare significance (-log(FDR)) and gene expression (log2FC) of reward- and region-specific DEG lists. Significant (p-value < 0.05) enriched KEGG gene sets were determined using g:Profiler (*30*) for each reward-specific seeking ensemble and were inputted into Cytoscape (*31*) for “EnrichmentMap” (*32*) network analysis to compare reward-specific gene sets.

## Results

### Cue-induced seeking of cocaine and sucrose recruit reward-specific ensembles

To characterize genetic profiles in cocaine- and sucrose-seeking ensembles, we used a dual self-administration (SA) and extinction (EXT) model and reward-specific cued-reinstatements, followed by fluorescently activated cell sorting (FACS) for downstream bulk RNA sequencing (RNAseq; Figure 1a). Ai14:cFos-TRAP2 mice underwent alternating daily sessions of intravenous cocaine and oral sucrose SA on a Fixed Ratio 1 (FR1) schedule (Figure 1b, Figure S1). SA was contingent on cue-paired interaction with cocaine- or sucrose-reinforced active nose pokes (NP). An additional non-reinforced inactive NP without consequences was also present. Cocaine (Figure S2a) and sucrose (Figure S2b) SA sessions showed consistent discrimination between the active NP and the inactive NP. Despite no sex differences observed during cocaine SA (Figure S3a), females interacted significantly more with the inactive NP than males during sucrose SA (Figure S3b). This finding is consistent with heightened cocaine preference during sucrose SA sessions for females when compared to males (Figure S3c). Overall, reward intake was equal for cocaine and sucrose and remained stable throughout SA (Figure S4a) with no sex differences (Figure S4b). Following SA, mice underwent EXT in the original SA chamber without reward administration or cue exposure. During EXT, mice decreased interaction and discrimination of NPs previously paired with cocaine or sucrose during SA (Figure 1c and Figure S5a) with no sex differences (Figure S5b,c).

**Figure 1.**
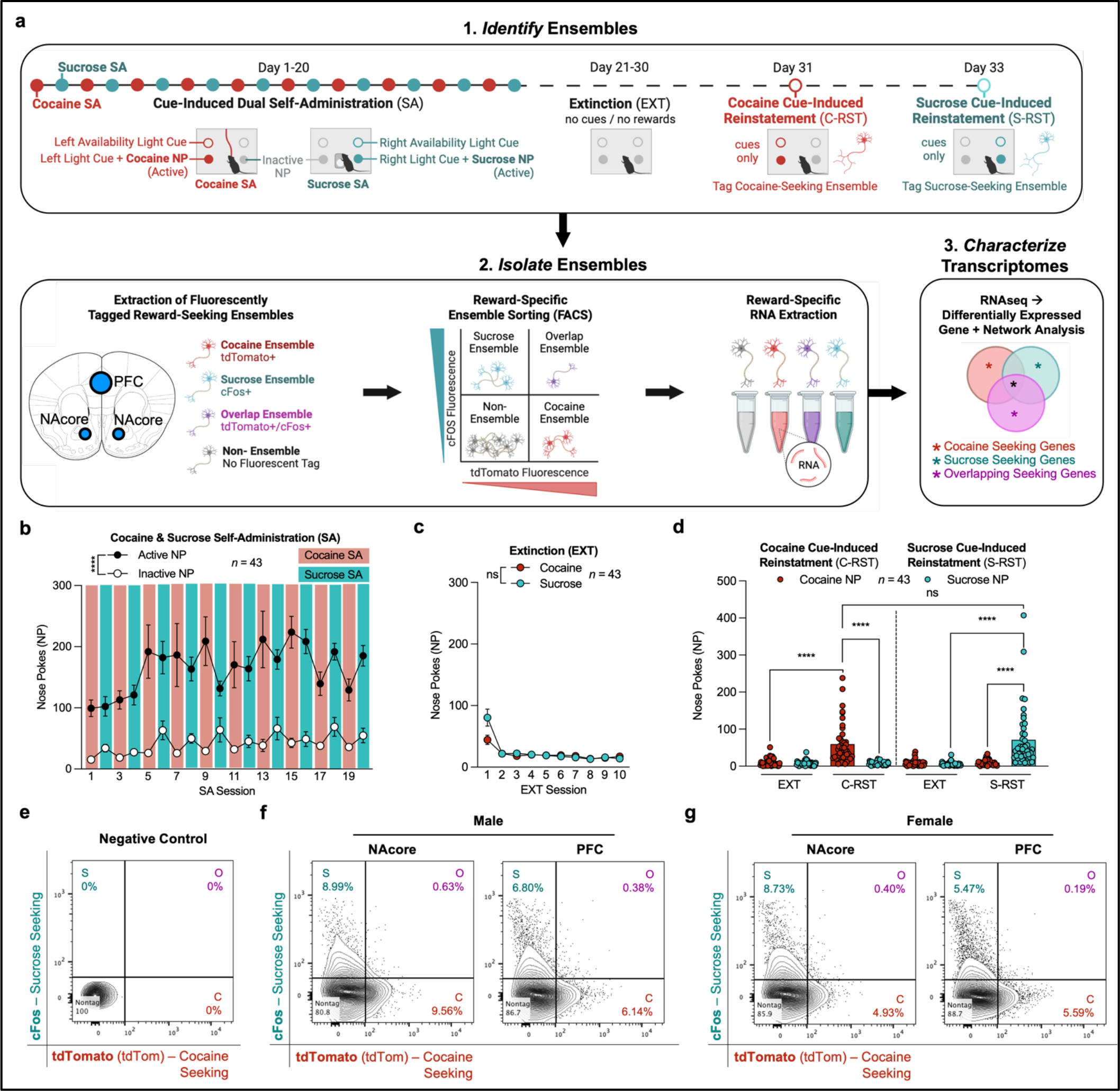
Polyreward self-administration of cocaine and sucrose and cell sorting of reward-specific ensembles. **a**. Behavioral training timeline for inducing cocaine and sucrose seeking for downstream fluorescently activated cell sorting and transcriptomic characterization of reward-specific ensembles from the nucleus accumbens core (NAcore) and prefrontal cortex (PFC) . **b**. During SA, mice interacted significantly more with the cocaine-seeking and sucrose-reinforced active NP (repeated measures two-way ANOVA for SA Active NP vs. SA Inactive NP over 20 sessions; NP F_(1, 84)_ = 70.85, P<0.0001; session F_(5.147, 432.3)_ = 3.343, P=0.0052; interaction F_(19, 1596)_ = 2.004, P=0.0063). **c**. During EXT, mice diminished differentiation between the active NP paired with cocaine or sucrose SA (repeated measures two-way ANOVA for EXT Cocaine NP vs. EXT Sucrose NP over 20 sessions; NP F_(1, 84)_ = 1.005, P=0.3190; session F_(1.758, 147.7)_ = 28.98, P<0.0001; interaction F_(9, 756)_ = 4.886, P<0.0001). **d**. Induction of cocaine-seeking and sucrose-seeking behavior displayed by significantly increased interaction with the active NP in comparison to the inactive NP during both C-RST (paired t-test for C-RST Cocaine NP vs. C-RST Sucrose NP; P<0.0001) and S-RST (paired t-test for S-RST Sucrose NP vs. S-RST Cocaine NP; P<0.0001) in addition to significantly increased interaction with the active NP in comparison to the level of active NP interaction in the last EXT session before C-RST (paired t-test for C-RST Cocaine NP vs. C-EXT Cocaine NP; P<0.0001) or S-RST (paired t-test for S-RST Sucrose NP vs. S-EXT Sucrose NP; P<0.0001). There was no difference when comparing levels of interaction with the active NP between C-RST and S-RST sessions (paired t-test for C-RST Cocaine NP vs. S-RST Sucrose NP; P=0.2399). **e**. Flow cytometric quadrant plot for negative control absent of antibody for cocaine-seeking and sucrose-seeking ensembles. **f,g**. Flow cytometric quadrant plots for male (**f**) and female (**g**) with percentages of cocaine-seeking neurons (C; bottom-right; tdTomato+/cFos-), sucrose-seeking neurons (S; top left; cFos+/tdTomato-) and overlapping neurons between cocaine-seeking and sucrose-seeking (O; top right: tdTomato+/cFos+) and non-ensemble (bottom-left; tdTomato-/cFos-) “nontag” neurons. Quadrant plots are a representative for male (Figure S2d) and female replicates (Figure S2e).

To induce seeking of cocaine and sucrose, mice underwent cocaine-(C-RST) and sucrose-specific (S-RST) cued-reinstatement sessions (Figure 1d, Figure S6), where mice were exposed to reward-specific cues in the absence of reward administration. Mice sought cocaine during the C-RST session, as demonstrated by increased interaction with the cocaine-paired active NP compared to the inactive NP and to the cocaine-paired active NP during the EXT session from the previous day (Figure 1d). Immediately following C-RST, 4-hydroxytamoxifen (4-OHT) was administered to tag active cFos-dependent neurons in the ensemble underlying cocaine-seeking behavior with fluorescent tdTomato. In parallel, mice sought sucrose during the S-RST session, as observed by increased interaction with the sucrose-paired active NP compared to the inactive NP and to the sucrose-paired active NP during the previous EXT session (Figure 1d, Figure S6). Mice were euthanized 60 minutes post-S-RST to capture endogenous cFos expression using immunocytochemistry (ICC) for neurons comprising sucrose-seeking ensembles while the tagged cocaine-seeking neuronal ensemble retained tdTomato expression. Despite differences in sex within SA, no sex differences were found in cue-induced cocaine-or sucrose-seeking behavior (Figure S6a).

Immediately following euthanasia, we dissociated neurons from the NAcore and PFC regions using FACS-separation of cocaine and sucrose-seeking neuronal ensembles. After debris removal (Figure S7a), we used an Ai14:cFos-TRAP2 control mouse with only a neuronal marker (NeuN+) and no ensemble tagging (Figure 1e, Figure S7b,c) to identify and sort cocaine-(C; lower right quadrant; tdTomato+) and sucrose-seeking (S; upper left quadrant; cFos+) neuronal ensembles in males (Figure 1f; Figure S7d) and females (Figure 1g; Figure S7e). We additionally sorted the overlapping ensemble (O; upper right quadrant; tdTomato+/cFos+), comprised of neurons activated during both cocaine and sucrose seeking. To ensure the downstream genetic analysis was specific to ensembles, we isolated neurons referred to as the non-ensemble (bottom left quadrant) that were not tagged in cocaine-seeking, sucrose-seeking, or overlapping ensembles. Additional flow cytometry analysis showed an overall significant main effect in the size of sex- and region-specific ensembles (Figure S8a-d). After identifying and isolating cocaine-seeking, sucrose-seeking, and overlapping ensembles, we proceeded with transcriptomic characterization.

### Cocaine and sucrose seeking feature reward-specific genetic alterations

To understand genetic regulation specific to cocaine or sucrose seeking, we sequenced the RNA of lysates from reward-specific seeking ensembles for significant DEG comparison (Supplemental Figure 9). Before we compared DEGs between reward-seeking ensembles, classical metric multidimensional scaling (MDS) of all samples revealed genetic profiles are defined by region-specific but not sex-specific variation (Supplemental Figure 10). Thus, we completed downstream DEG comparisons in a region-specific manner including sex as a variable (Supplemental Figure 11,12). DEG data sets for cocaine-seeking, sucrose-seeking, and overlap ensembles, in addition to non-ensemble neurons, were determined in reference to naïve data sets (Supplemental Figure 11,12; Supplemental Table 1,5).

Comparing significant reward-specific seeking DEGs in neuronal ensembles from the NAcore (Figure 2a-d) resulted in 310 cocaine-seeking, 1997 sucrose-seeking, and 516 overlapping ensemble-specific DEGs excluding DEGs related to the non-ensemble neuronal population (Figure 2e). The extremities of gene expression, i.e., the ten highest expressed and ten lowest expressed DEGs for each reward specific ensemble in the NAcore, based on log transformation of the fold change (log2FC), showed robust differential expression in comparison to the expression within the other ensembles (Figure 2g, Table 2). Dopaminergic gene *Foxa2* (*33*) and serotonergic gene *Fev* (*34*) had higher expression in the NAcore cocaine-seeking ensemble compared to the NAcore sucrose-seeking and overlapping ensembles. Glucose-regulating genes were upregulated (*Dgat1*) (*35, 36*), or downregulated (*Ttc4*) (*37*), in the NAcore sucrose-seeking ensemble compared to the NAcore cocaine-seeking and overlapping ensembles (Figure 2h, Table 3). Additionally, glucose-regulating *Ube2i* (*37*) expression was downregulated for the NAcore overlapping ensemble in comparison to the NAcore cocaine-seeking and sucrose-seeking ensembles (Figure 2i, Table 4).

**Figure 2.**
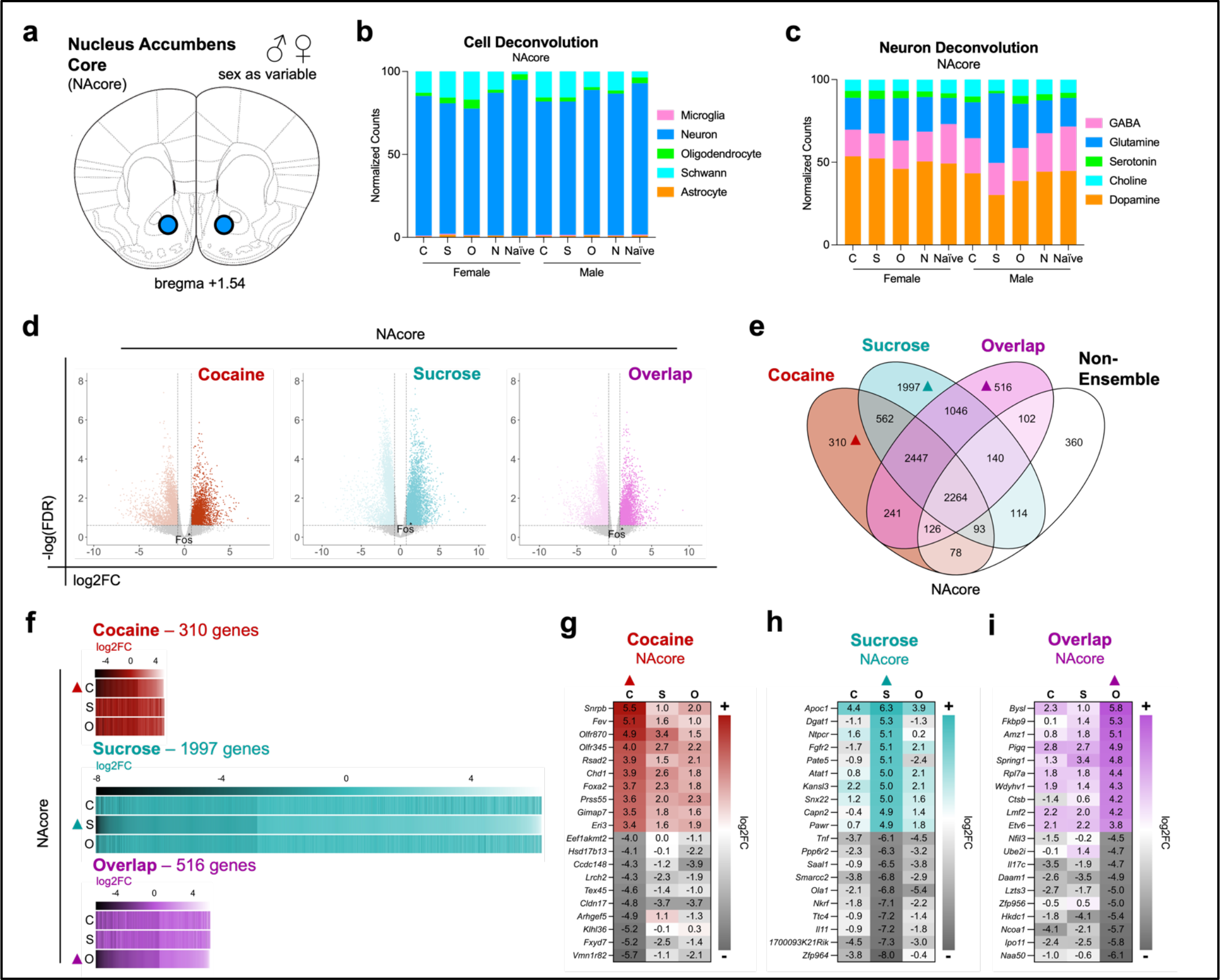
Cocaine-seeking and sucrose-seeking ensembles have overlapping and discrete genetic factors within the nucleus accumbens core (NAcore). **a**. Schematic of NAcore region for RNA extraction. **b**. Cell deconvolution for NAcore samples (cocaine-seeking ensemble = C; sucrose-seeking ensemble= S; overlapping ensemble= O; non-ensemble = N) to determine percent of neurons (*Rbfox3*/NeuN), microglia (*Aif1*/Iba1), oligodendrocytes (*Olig1*/Olig1), Schwann (*Ngfr*/p75ntr), and astrocyte (*s100b*/S100b) cell types. **c**. Neuron cell-type deconvolution for NAcore samples to determine percent of dopaminergic (*Drd1, Drd2, Foxa2*), GABAergic (*Gad1, Slc6a1, Gabbr1*), glutaminergic (*Glul, Grin1, Slc17a7*), serotoninergic (*Ptprc, Slc6a4, Fev*), cholinergic (*Ache, Chat*) cell types. **d**. Volcano plots representing reward-specific DEGs scaled by -log_10_(FDR) (y-axis; factor representing significance) and log_2_(fold-change) (log2FC; x-axis; factor representing gene expression) within the NAcore. **e**. Four-way Venn diagrams comparing differentially expressed genes (DEG; in reference to naïve mice) in cocaine-seeking, sucrose-seeking, and overlapping ensembles in addition to the non-ensemble population within the NAcore. Upside down triangles denote the NAcore DEGs defined as cocaine-seeking (red triangle), sucrose-seeking (aqua triangle), or overlapping (purple triangle) for downstream characterization. **f**. Heatmap of DEGs specific to cocaine-seeking, sucrose-seeking, and overlapping ensembles in the NAcore. Colored triangles denote the NAcore DEGs (from comparison in Figure 2e) used for comparison with log2FC of the same gene in other reward-seeking ensembles. **g,h,i**. Heatmap for the ten highest and ten lowest expressed DEGs specific to cocaine-seeking (**g**), sucrose-seeking (**h**), and overlapping ensembles (**i**) in the NAcore. Colored triangles denote the NAcore DEGs (from comparison in Figure 2e) used for comparison with log2FC of the same gene in other reward-seeking ensembles.

Comparing significant reward-specific seeking DEGs in neuronal ensembles from the PFC (Figure 3a-d) resulted in 552 cocaine-seeking, 856 sucrose-seeking, and 494 overlapping ensemble-specific DEGs excluding DEGs related to the non-ensemble neuronal population (Figure 3e). Similar to the NAcore, extremities of reward specific DEGs in the PFC displayed robust specificity of differential expression compared to the expression of the same genes in the other ensembles. For instance, glycoprotein-coding *Scube1* (*38, 39*) was downregulated in the PFC cocaine-seeking ensemble compared to the PFC sucrose-seeking and overlapping ensembles (Figure 3g, Table 6). Similar to the NAcore, glucose-regulating genes were upregulated (*Gng12*) (*37*), or downregulated (*Pmm1*) (*40, 41*), in the PFC sucrose-seeking ensemble compared to the PFC cocaine-seeking and overlapping ensembles (Figure 3h, Table 7). In addition, glucose-regulating *Hecw1* (*37, 42*) expression was downregulated for the PFC overlapping ensemble in comparison to the PFC cocaine-seeking and sucrose-seeking ensembles (Figure 3i, Table 8). Together, these data show that genetic expression underlying cocaine and sucrose seeking is both shared and distinct in the NAcore or the PFC.

**Figure 3.**
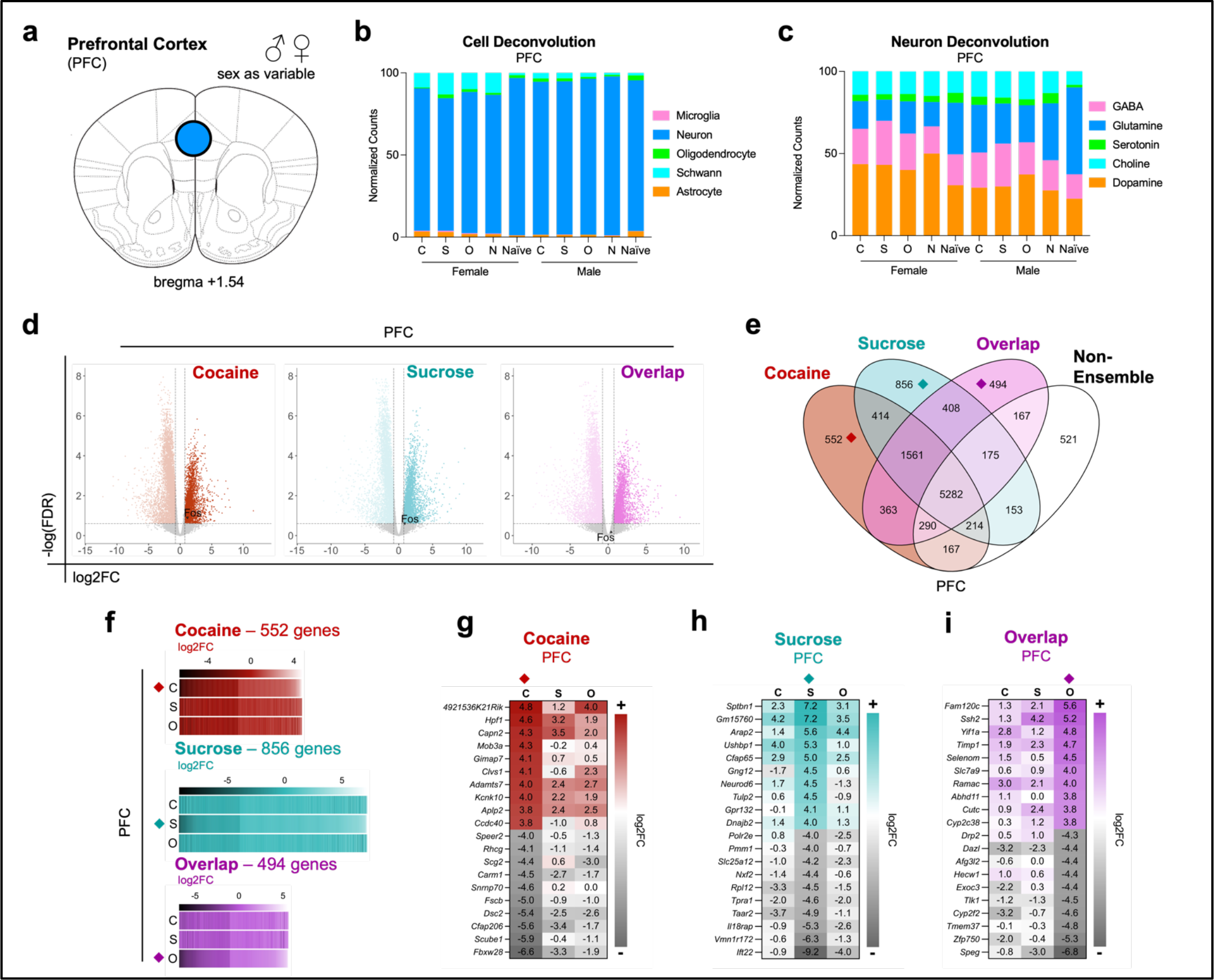
Cocaine-seeking and sucrose-seeking ensembles have overlapping and discrete genetic factors within the prefrontal cortex (PFC). **a**. Schematic of PFC region for RNA extraction. **b**. Cell deconvolution for PFC samples (cocaine-seeking ensemble = C; sucrose-seeking ensemble= S; overlapping ensemble= O; non-ensemble = N). **b**. Cell deconvolution for PFC samples (cocaine-seeking ensemble = C; sucrose-seeking ensemble= S; overlapping ensemble= O; non-ensemble = N) to determine percent of neurons (*Rbfox3*/NeuN), microglia (*Aif1*/Iba1), oligodendrocytes (*Olig1*/Olig1), Schwann (*Ngfr*/p75ntr), and astrocyte (*s100b*/S100b) cell types. **c**. Neuron cell-type deconvolution for PFC samples to determine percent of dopaminergic (*Drd1, Drd2, Foxa2*), GABAergic (*Gad1, Slc6a1, Gabbr1*), glutaminergic (*Glul, Grin1, Slc17a7*), serotoninergic (*Ptprc, Slc6a4, Fev*), cholinergic (*Ache, Chat*) cell types. **e**. Four-way Venn diagrams comparing differentially expressed genes (DEG; in reference to naïve mice) in cocaine-seeking, sucrose-seeking, and overlapping ensembles in addition to the non-ensemble population within the PFC. Upside down diamonds denote the PFC DEGs defined as cocaine-seeking (red diamond), sucrose-seeking (aqua diamond), or overlapping (purple diamond) for downstream characterization. **f**. Heatmap of DEGs specific to cocaine-seeking, sucrose-seeking, and overlapping ensembles in the PFC. Colored diamonds denote the PFC DEGs (from comparison in Figure 3e) used for comparison with log2FC of the same gene in other reward-seeking ensembles. **g,h,i**. Heatmap for the ten highest and ten lowest expressed DEGs specific to cocaine-seeking (**g**), sucrose-seeking (**h**), and overlapping ensembles (**i**) in the PFC. Colored diamonds denote the PFC DEGs (from comparison in Figure 3e) used for comparison with log2FC of the same gene in other reward-seeking ensembles.

### Cocaine and sucrose seeking have dynamic genetic signatures regulating neuronal signaling

To further characterize cocaine- and sucrose-seeking DEGs, we utilized gene sets within the Kyota Encyclopedia of Genes and Genomes (KEGG) database to identify genetic networks within cocaine-seeking, sucrose-seeking, and overlapping ensembles. For this, we used previously discussed DEGs specific for each reward-seeking ensemble in the NAcore (Figure 2e) and PFC (Figure 3e). Analysis of enriched KEGG pathways for each reward-specific seeking ensemble in the NAcore resulted in 6 cocaine-seeking pathways, 29 sucrose-seeking pathways, 9 overlapping pathways, and 4 pathways shared by one or more ensembles (Figure 4a). We examined gene-level expression profiles for the “dopaminergic synapse” pathway enriched in the NAcore cocaine-seeking ensemble (Figure 4b) given the importance of dopamine regulation in reward-related behavior (*43*). Within this pathway, some genes had discrete expression for cocaine-seeking, sucrose-seeking, or overlapping ensembles.

**Figure 4.**
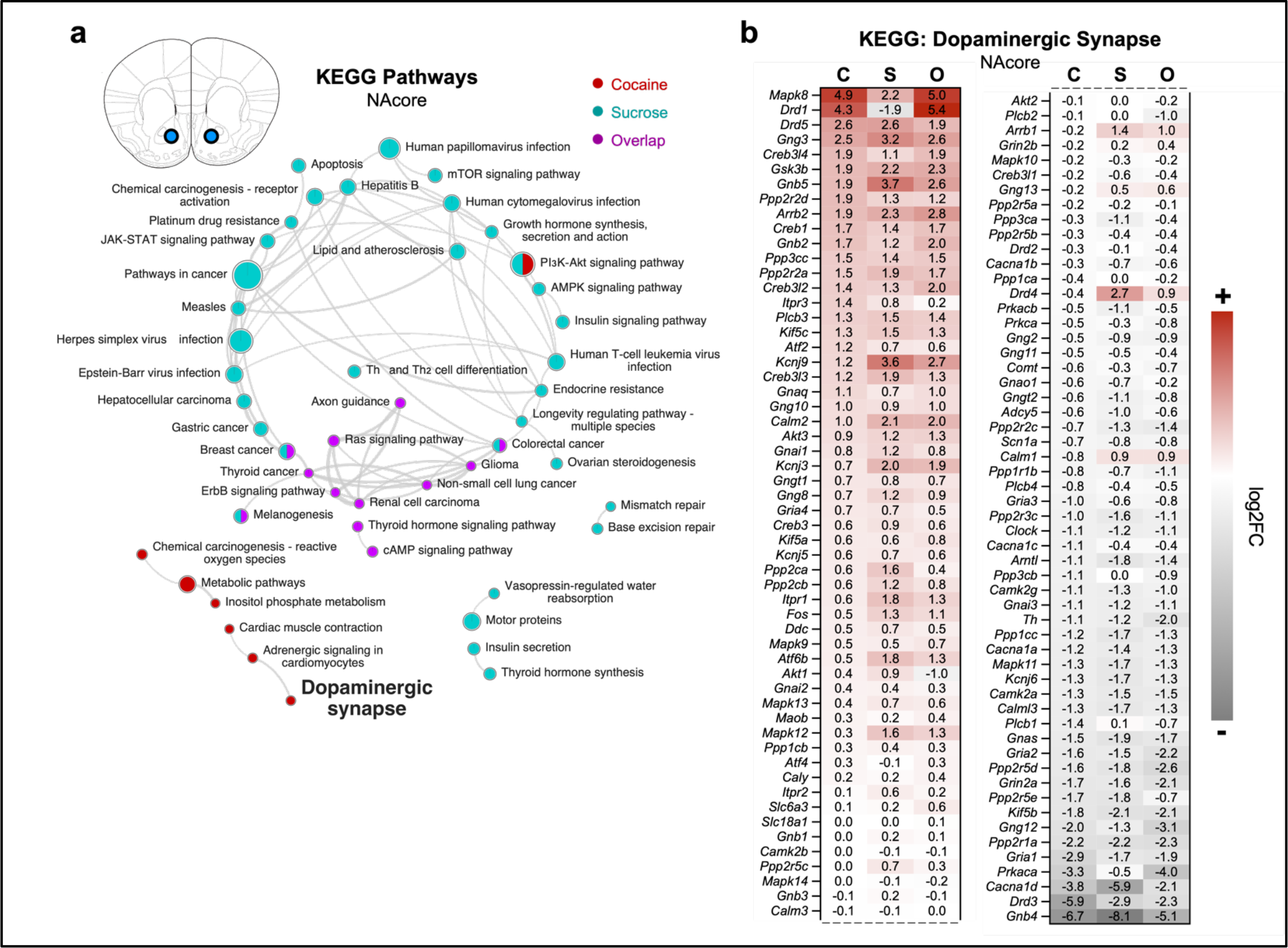
Cocaine-seeking and sucrose-seeking ensembles have similar and discrete genetic ontology within the nucleus accumbens core (NAcore). **a**. Network representation for KEGG gene set enrichment analysis (g:Profiler) of reward-specific DEGs in the NAcore (from comparison in 2e). Colored circular nodes represent enriched KEGG pathways colored by reward (cocaine-seeking ensembles = red; sucrose-seeking ensembles = aqua; overlapping ensembles = purple; shared by multiple types of reward seeking ensembles = multicolored fractions). Lines connecting circular nodes represent similarity of genes within two KEGG pathways. **b**. Heatmap of all DEGs annotated in the KEGG “dopaminergic synapse” pathway enriched in cocaine-seeking in the NAcore from Figure 4a with log2FC comparisons of genes in all reward-seeking ensembles (cocaine-seeking ensemble = C; sucrose-seeking ensemble= S; overlapping ensemble= O).

The family of dopaminergic receptors (*44*) saw dynamic expression: *Drd3* was downregulated in all ensembles with lowest expression in cocaine-seeking while *Drd5* was upregulated for all seeking ensembles with similar magnitude, *Drd2* was slightly under-expressed in each seeking ensemble, *Drd1* was upregulated downregulated in sucrose seeking while upregulated in both the cocaine-seeking and overlapping ensemble, *Drd4* was downregulated in cocaine-seeking compared to sucrose-seeking and overlapping ensembles. In addition to dopaminergic receptors, dopamine production via tyrosine hydroxylase (*Th*) expression (*45*) was downregulated for all seeking ensembles with the lowest expression in the overlapping ensemble. We additionally observed dynamic expression of *Creb* isoforms (*46, 47*) with upregulation of *Creb1, Creb3, Creb3l2, Creb3l3*, and *Creb314* in contrast to slight downregulation of *Creb3l1* for all seeking ensembles. Gene expression for cocaine-linked potassium channels (*Kcnj3* and *Kcnj9*) (*48, 49*) was the lowest for cocaine-seeking compared with sucrose-seeking and overlapping ensembles. Cocaine-linked calcium channel gene expression (*Cacna1c* and *Cacna1d*) (*50–52*) was downregulated overall for every seeking ensemble with the lowest expression for *Cacna1c* in cocaine-seeking while *Cacna1d* had the lowest expression in sucrose-seeking.

Analysis of enriched KEGG pathways for each reward-specific seeking ensemble in the PFC resulted in 22 cocaine-seeking pathways, 6 sucrose-seeking pathways, 9 overlapping pathways, and 3 pathways shared by one or more ensembles (Figure 5a). We saw enrichment of the “cocaine addiction” pathway for cocaine-seeking ensembles. Some genes within this pathway had differing expressions between cocaine-seeking, sucrose-seeking, and the overlapping ensembles.

**Figure 5.**
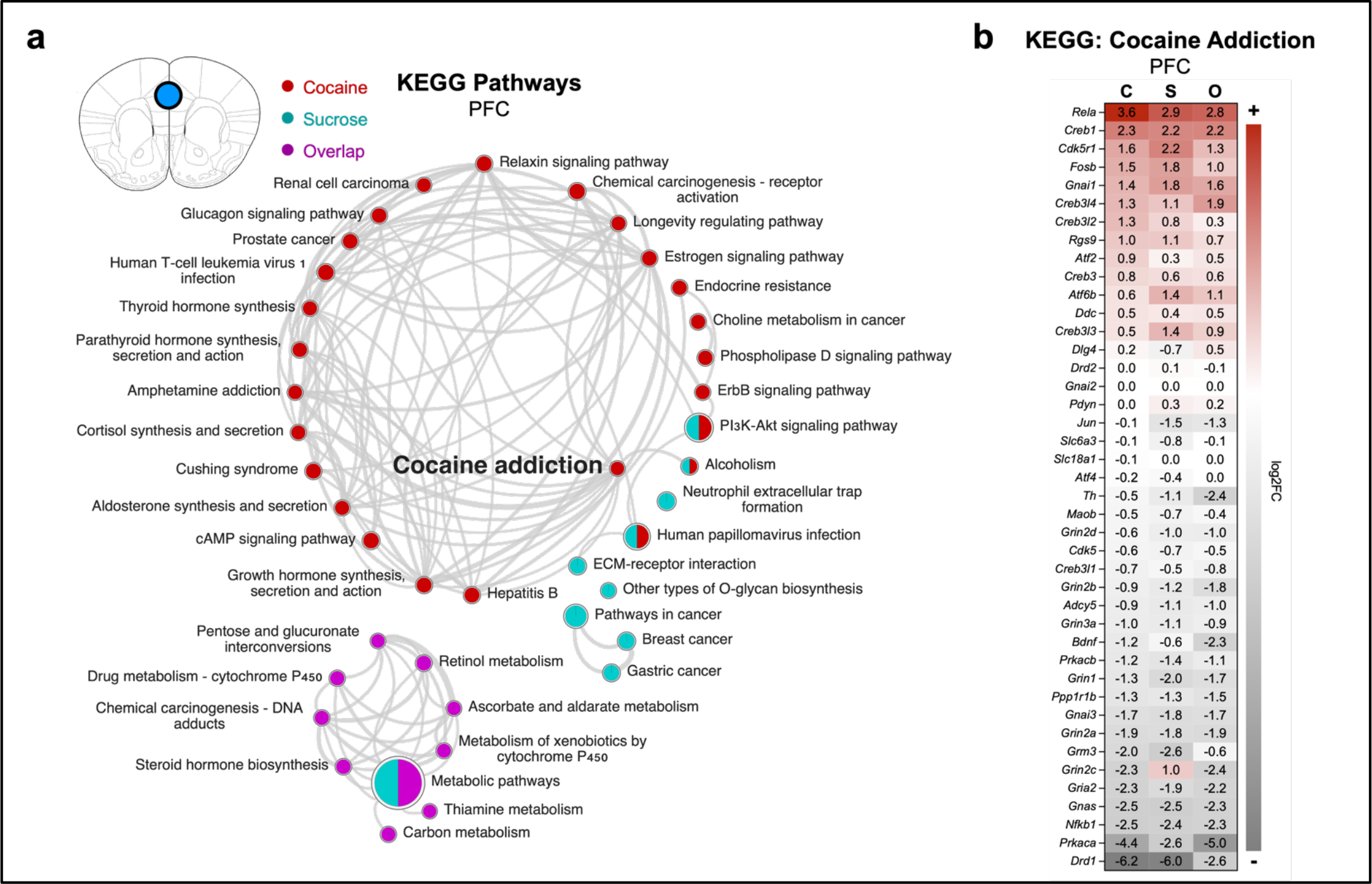
Cocaine-seeking and sucrose-seeking ensembles have similar and discrete genetic ontology within the prefrontal cortex (PFC). **a**. Network representation for KEGG gene set enrichment analysis (g:Profiler) of reward-specific DEGs in the PFC (from comparison in 3e). Colored circular nodes represent enriched KEGG pathways colored by reward (cocaine-seeking ensembles = red; sucrose-seeking ensembles = aqua; overlapping ensembles = purple; shared by multiple types of reward seeking ensembles = multicolored fractions). Lines connecting circular nodes represent similarity of genes within two KEGG pathways. **b**. Heatmap of all DEGs annotated in the KEGG “cocaine addiction” pathway enriched in cocaine-seeking in the PFC from Figure 5a with log2FC comparisons of genes in all reward-seeking ensembles (cocaine-seeking ensemble = C; sucrose-seeking ensemble= S; overlapping ensemble= O).

We saw contrasting under expression for *Drd1* in the PFC cocaine-seeking and overlapping ensembles in comparison to the overexpression of NAcore expression, while sucrose-seeking displayed further downregulation in the PFC. *Drd2* had a minimal fluctuation in the sucrose-seeking and overlapping ensembles and no expression change within the cocaine-seeking ensemble. Similar to the NAcore, dopamine regulating *Th* was under-expressed for each seeking ensemble; however, the findings were more dynamic in the PFC with the three-fold lower expression in the overlapping ensemble compared to the cocaine-seeking ensemble. Aligning with our results from the NAcore, the PFC featured upregulation of *Creb1, Creb3, Creb3l2, Creb3l3*, and *Creb314* in contrast to slight downregulation of *Creb3l1* for all seeking ensembles. Additionally, under expression for genetic subunits for cocaine-linked NMDAR signaling (*Grin2a, Grin2b, Grin2c, Grin2d*, and *Grin3a*) (*53, 54*) was observed for each seeking ensemble with the exception for *Grin2c* that was only upregulated in the sucrose-seeking ensemble. We also found key transcription factors linked with cocaine seeking (*55*) to be upregulated the highest in cocaine-seeking compared to sucrose seeking (*Atf2*), upregulated for all ensembles similarly (*Fosb*), or downregulated more than ten-fold in the sucrose-seeking and overlapping ensemble compared to cocaine-seeking (*Jun*). Dynamic *Bdnf* expression, associated with cocaine sensitization (*56*), was downregulated in all seeking ensembles with the lowest expression in the overlapping ensemble.

Building on genes previously mentioned in the NAcore sucrose-seeking ensemble, we found “insulin signaling pathways” enriched only in the sucrose-seeking ensemble. We also found pathways correlating with previous animal studies focusing on cocaine and sucrose exposure enriched for the overlapping ensemble in the NAcore, e.g., “cAMP signaling pathway” (56, 57), or PFC, e.g., “steroid hormone biosynthesis” (*57, 58*). Additionally, we saw shared enrichment of the “alcoholism” pathway for both cocaine and sucrose-seeking ensembles supporting claims that there is common liability between addictive substances (*39*). The “PI3K-Akt signaling pathway” was enriched for sucrose- and cocaine-seeking ensembles in both the NAcore and PFC, correlating with previous studies (*56, 59*). Taken together, these data highlight genetic pathways underlying cocaine and sucrose seeking are both shared and distinct in the NAcore or the PFC.

### Cocaine and sucrose seeking have region-dependent genetic signatures between the NAcore and PFC

We further compared reward-specific DEGs between the NAcore (Figure 2e) and PFC (Figure 3e) to determine region-dependent alterations within cocaine and sucrose-seeking ensembles. Although most genes were region-specific, we observed that genes shared between regions for each reward-seeking ensemble had a pattern for discrete gene expression (Figure 6a-d). Within the 12 DEGs shared between the NAcore and PFC in cocaine-seeking ensembles, we found that GTP-binding-protein-coding *Gimap7* (*60*) was upregulated in the NAcore and PFC in cocaine-seeking ensembles (Figure 6e; Table 9) in comparison to the sucrose-seeking and overlapping ensembles with more pronounced upregulation of *Gimap7* in the PFC (Figure 6e). Furthermore, we found 247 genes shared between the NAcore and PFC sucrose-seeking ensembles with mostly unique genetic expression compared to cocaine-seeking and overlapping ensembles (Figure 6f,g; Table 10). Dopamine-regulating *Pgam5* (*61, 62*) was under-expressed in the NAcore and overexpressed in the PFC for sucrose-seeking ensembles compared to cocaine-seeking and overlapping ensembles. Additionally, we found discrete expression patterns within the 30 genes associated with the overlapping ensemble (Figure 6h; Table 11). For example, DNA-repair-regulating *Prkdc* (*63, 64*) was downregulated in both the NAcore and PFC for the overlapping ensemble compared to cocaine-seeking and sucrose-seeking ensembles, with more pronounced downregulation in the PFC.

**Figure 6.**
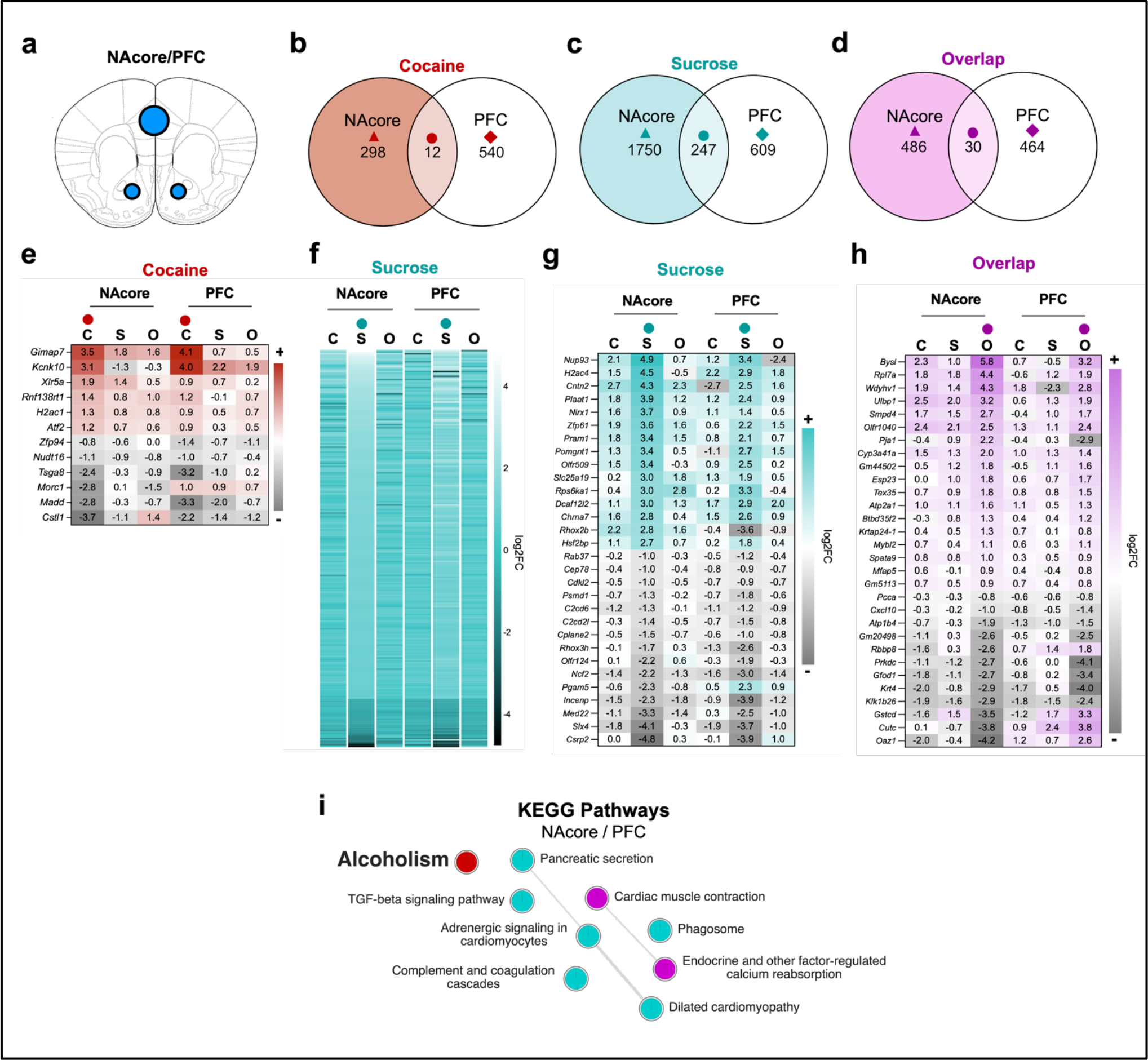
Cocaine-seeking and sucrose-seeking ensembles have a similar and discrete genetic expression in the nucleus accumbens core (NAcore) and prefrontal cortex (PFC). **a**. Schematic of NAcore and PFC regions. **b,c,d**. Venn diagrams to compare reward-specific DEGs between the NAcore and PFC for cocaine-seeking (**b**), sucrose-seeking (**c**), and overlapping (**d**) ensembles. Colored triangles represent reward-specific DEGs from comparison 2e for the NAcore. Colored circles denote the shared NAcore-PFC DEGs in cocaine-seeking (red circle), sucrose-seeking (aqua circle), or overlapping (purple circle) ensembles for downstream characterization. **e,f,g,h**. Heatmap for DEGs specific to cocaine-seeking (**e**), sucrose-seeking (**f**), ten highest and ten lowest expressed DEGs in sucrose-seeking (**g**), and overlapping ensembles (**h**) in the PFC. Colored circles denote the reward-specific shared NAcore-PFC DEGs (from comparison in Figure 6b-d) used for comparison with log2FC of the same gene in other reward-seeking ensembles in the NAcore and PFC. **i**. Network representation for KEGG gene set enrichment analysis (g:Profiler) of reward-specific DEGs shared between the NAcore and PFC (from comparison in Figure 6b-d). Colored circular nodes represent enriched KEGG pathways colored by reward (cocaine-seeking ensembles = red; sucrose-seeking ensembles = aqua; overlapping ensembles = purple; shared by multiple types of reward seeking ensembles = multicolored fractions). Lines connecting circular nodes represent similarity of genes within two KEGG pathways.

To further characterize cocaine- and sucrose-seeking DEGs shared between both the NAcore and PFC, we again used the KEGG database to identify genetic networks within cocaine-seeking, sucrose-seeking, and overlapping ensembles (Figure 6i). Mentioned previously to have enrichment for both cocaine and sucrose seeking in the PFC, the “alcoholism” pathway was only enriched for cocaine-seeking when shared between the NAcore and PFC. We saw enrichment of the “pancreatic secretion” pathway in sucrose-seeking for both the NAcore and PFC, aligning with our previously mentioned glucose-regulating gene expression findings and other studies relating to the gut-brain axis (*65*). Additionally, enrichment of the “endocrine and other factor-regulated calcium reabsorption” supports previous studies relating calcium dysregulation to cocaine- and sucrose-exposed animals (*66–68*). To summarize, genetic alterations and associated pathways linked to cocaine- and sucrose-seeking ensembles are region-dependent.

## Discussion

We used within-subject cocaine and sucrose dual SA conditioning followed by extinction training and cued reinstatement to induce cocaine and sucrose seeking in cFos-TRAP2 mice (Figure 1b-d). Through FACS (Figure 1e-g), we isolated neurons from reward-specific seeking ensembles and the non-ensemble population for downstream genetic profiling compared to naïve controls. Within two critical reward-signaling regions, the NAcore and PFC, we used an unbiased approach to identify genetic signatures exclusive to or shared between cocaine- and sucrose-seeking ensembles in addition to ensembles activated during both cocaine and sucrose seeking, i.e., overlapping ensemble.

### Cocaine- and sucrose-seeking ensembles feature unique and shared genetic expression linked to neurotransmitter regulation and glucose responses

Compared to naïve mice not exposed to rewards and non-ensemble cells in mice exposed to cocaine and sucrose, we identified significant ensemble-specific genetic alterations in cocaine-seeking, sucrose-seeking, and overlapping ensembles in the NAcore (Figure 2e) and PFC (Figure 3e). Observing the extremities of reward-specific gene expression in the NAcore (Figure 2g-i) and PFC (Figure 3g-i) revealed patterns of exclusive or similar gene expression for cocaine-seeking, sucrose-seeking, or overlapping ensembles for both sexes. We further identified unique and overlapping genetic pathways enriched for reward-seeking ensembles in the NAcore (Figure 4) and PFC (Figure 5).

Within the NAcore, we found dynamic expression of neurotransmitter regulatory genes in cocaine- and sucrose-seeking ensembles. Specifically, we saw a dynamic expression of genes regulating dopamine production and receptor signaling amongst all reward-seeking ensembles, with upregulation of *Foxa2* in the cocaine-seeking ensemble and downregulation of *Drd1* in the sucrose-seeking ensemble. Further, we found upregulated expression of serotoninergic *Fev* in cocaine-seeking, which has previously been linked with facilitating cocaine preference via *Drd1* neurons (*34*). Interestingly, we found enrichment of the “dopaminergic synapse” pathway only in the cocaine-seeking ensemble. Here, we’ve highlighted the genetic complexity of neurotransmitter regulation involved in both cocaine and sucrose seeking. Additionally, we found expression of genes regulating reinforcement of sucrose-seeking and glucose regulation linked to the sucrose-seeking and overlapping ensembles in the NAcore. These findings align with the catabolism of sucrose into glucose and fructose (*69*). Specifically, *Ttc4* in the sucrose-seeking ensemble and *Ube2i* in the overlapping ensemble were considerably under-expressed and have previously been linked with regulation of the glycolytic enzyme glucokinase (Gck) in paraventricular thalamus (PVT) neurons (*37*). Going further, glucose-responsive Gck-dependent neural circuits between the nucleus accumbens and PVT have been linked to overconsumption of sucrose. We also report overall enrichment for insulin signaling in the sucrose-seeking ensemble (Figure 4). Interestingly, previous studies have highlighted the neural connections between the gut and the PVT, with receptors featured in both regions to facilitate crosstalk signaling (*70*).

Like in the NAcore, we observed distinct or shared genetic expression within reward-seeking ensembles in the PFC, including genes differentially expressed regulating neurotransmitter production or signaling. Opposite of the NAcore, considerable downregulation of *Drd1* occurred in the PFC, suggesting region-specific regulation of dopamine signaling during both cocaine and sucrose seeking. In the PFC cocaine-seeking ensemble, we found downregulation of *Scube1*, which was identified as having single nucleotide polymorphisms linked with the common liability theory of substance use disorder, according to which there are underlying genes shared to facilitate addictive-like phenotypes for any substance (*39*). Supporting our hypothesis that sorted populations are specific to cocaine or sucrose seeking, we found enrichment of the “cocaine addiction” pathway in only the cocaine-seeking ensemble, which highlighted unique gene expression for *Jun, Th, Bdnf*, and *Atf2*. In parallel to the dynamic expression of glucose-regulating genes in the NAcore, we also found genes in the PFC regulating glucose responses. For instance, *Gng12* (upregulated in the sucrose-seeking ensemble) and *Hecw1* (downregulated in the overlapping ensemble) have also been previously linked with the regulation of Gck in PVT neurons (*37*). *Hecw1* has also been linked with overdose of cocaine in post-mortem human tissue (*42*). We also report an overlap of the “alcoholism” pathway enriched in the cocaine-seeking ensemble in both regions (Figure 5).

Going further, we saw both discrete or shared expression of DEGs shared between the NAcore and PFC (Figure 6). GTP-binding protein *Gimap7*, shown to be enriched after methamphetamine exposure in a non-human primate study (*60*), was overexpressed in both regions in the cocaine-seeking ensemble. In addition, we again saw genes related to glucose responses during obesity resistance with discrete expression in the sucrose-seeking ensemble (*Pgam5*) or the overlapping ensemble (*Prkdc*), which is interesting considering the glucose-linked satiety effects linked with chronic cocaine exposure (*69*). Specifically, the expression of glucose regulating *Pgam5* has previously been shown to alter dopaminergic signaling. We also found enrichment of “pancreatic secretion” in the sucrose-seeking ensemble. Notably, the cocaine-seeking ensemble does not recruit glucose-responsive genes or pathways, which we find are only featured in ensembles related to sucrose seeking. Overall, these results suggest that, in the NAcore and PFC, cocaine- and sucrose-seeking ensembles have shared and discrete genetic signatures with an additional unique gene signature tied to the overlapping ensemble. For future studies, differentially expressed genes exclusive to cocaine-seeking ensembles should be prioritized, i.e., manipulating target genes in the NAcore or PFC and observing the effect on cocaine- and sucrose-seeking behaviors.

### Cell-type specificity within reward-specific ensembles and bioinformatics analysis

While it is known that different cell types are involved in ensembles (*71*), we limited our FACS procedure to isolate neuronal ensembles exclusively. From our flow cytometric results, we determined the size of female and male cocaine- and sucrose-seeking ensembles are equivalent in the NAcore and PFC except for the female PFC where the sucrose-seeking ensemble was larger (Figure S8a-d). We suggest the difference in ensemble size could be due to variability in the tagging process or possibly linked to the estrous cycle, discussed further. Considering the size differences in ensembles, we note that an equal number of events were sorted for each reward-seeking ensemble for technical normalization. We further conducted normalization of RNAseq counts within the edgeR/Limma pipeline to address inconsistencies within the sorting process (Supplemental Figure 10-12). We also highlighted similarity of cell-type populations (Figure 2b-c, Figure 3b-c) between FACS-sorted cell populations being compared downstream. The Limma pipeline used here determined DEGs for each reward-seeking ensemble compared to the naïve reference samples using adjusted p-values (FDR < 0.25) (*20, 72*) and a log2FC cutoff (>0.75; <-0.75). It should be noted that heatmaps feature comparisons of reward-specific genes with expression in other ensembles that did not undergo FDR and log2FC filtering. Full lists with FDR are included in these analyses (Supplemental Table 1-11).

### Sex differences are defined in cocaine- and sucrose-seeking behavior

We identified considerable behavioral sex differences supporting previous findings of sex-dependent characteristics in CUD (*73*). While we did not observe a higher sucrose intake in females shown in a previous study (8), females compared to males interacted significantly more with the NP associated with cocaine during the sucrose sessions (Figure S3b) and showed higher cocaine preference during sucrose self-administration (Figure S3c). Females could be more susceptible to the maladaptive rewarding effects of cocaine or could show learning delays after cocaine exposure compared to males. Cue-induced seeking behavior was equal in male and female mice (Figure S1l), which is different from what is seen in female outpatients, who are more likely to report increased craving in response to cocaine-associated cues than males (*74*). It should be noted we did not record the estrous cycle in this study to avoid stress-induced responses (*75*), but it is known that the estrous cycle influences cocaine-seeking (*76*) and food rewards (*77*) in female rats. Moreover, the multidimensional scaling (MDS) analysis of all samples did not reveal an effect of sex in genetic expression. Thus, we completed downstream DEG comparisons in a region-specific manner.

## Conclusion

We identified, isolated, and characterized specific and shared patterns of gene expression within cocaine-seeking, sucrose-seeking, and overlapping ensembles. We ultimately found a myriad of discrete or shared genetic signatures between each cocaine and sucrose seeking linked to neurotransmitter and glucose regulation. This repository of reward-specific genetic information is a valuable resource for further functional validation of target genes to reduce cocaine-seeking without altering natural reward-seeking behavior.

## Supporting information

Supplementary Figues and Legends

## Acknowledgments

The authors thank the University of Colorado Anschutz Genomics and Microarray Core for performing the RNA sequencing and all the members of the Bobadilla Lab for their help and support.

