## Supplementary Figues and Legends for "Differential genetic expression within reward-specific ensembles in mice"

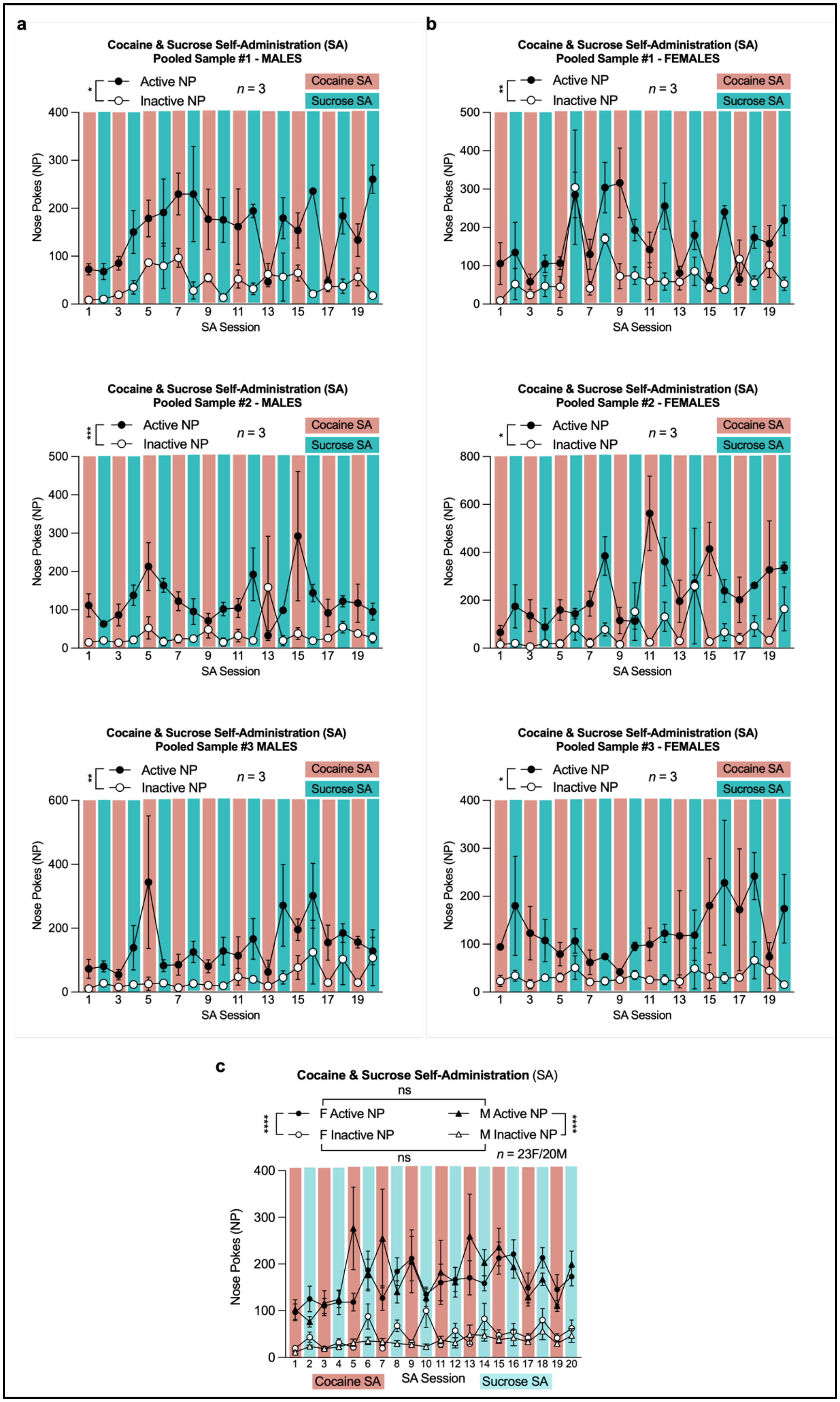


**Supplemental Figure 1 | Self-administration behavior of RNA-sequenced pools and sex-specific behavior. a.** Pooled male sample self-administration plots grouped by RNA pooling schematic of 3 mice per sample per three replicates. All male pooled samples displayed significant differentiation between the active NP and inactive NP regardless of reward throughout self-administration (repeated measures two-way ANOVA for RNA Male Pooled Sample 1: SA Active NP vs. SA Inactive NP over 20 sessions; session F (2.414, 9.657) = 2.912, P=0.0963; NP F (1, 4) = 20.17, P=0.0109; interaction F (19, 76) = 2.242, P=0.0071; repeated measures two-way ANOVA for RNA Male

Pooled Sample 2: SA Active NP vs. SA Inactive NP over 20 sessions; session F (2.358, 9.432) = 0.9774, P=0.4249; NP F (1, 4) = 86.52, P=0.0007; interaction F (19, 76) = 1.489, P=0.1138; repeated measures two-way ANOVA for RNA Male Pooled Sample 3: SA Active NP vs. SA Inactive NP over 20 sessions; session F (2.172, 8.689) = 1.635, P=0.2503; NP F (1, 4) = 37.08, P=0.0037; interaction F (19, 76) = 0.7943, P=0.7065). **b.** Pooled female sample self-administration plots grouped by RNA pooling schematic of 3 mice per sample per three replicates. All female pooled samples displayed significant differentiation between the active NP and inactive NP regardless of reward throughout self-administration (repeated measures two-way ANOVA for RNA Female Pooled Sample 1: SA Active NP vs. SA Inactive NP over 20 sessions; session F (3.238, 12.95) = 4.061, P=0.0288; NP F (1, 4) = 59.39, P=0.0015; interaction F (19, 76) = 1.400, P=0.1528); repeated measures two-way ANOVA for RNA Female Pooled Sample 2: SA Active NP vs. SA Inactive NP over 20 sessions; session F (2.436, 9.746) = 2.293, P=0.1477; NP F (1, 4) = 15.14, P=0.0177; interaction F (19, 76) = 1.679, P=0.0591); repeated measures two-way ANOVA for RNA Female Pooled Sample 3: SA Active NP vs. SA Inactive NP over 20 sessions; session F (2.239, 8.956) = 0.9729, P=0.4242; NP F (1, 4) = 8.744, P=0.0417; interaction F (19, 76) = 0.7388, P=0.7677). **c.** Female (F) and male (M) sex-specific plots for cocaine and sucrose self-administration for females (repeated measures two-way ANOVA for F SA Active NP vs. F SA Inactive NP over 20 sessions; NP $F_{(1, 44)}$ = 46.21, P<0.0001; session $F_{(7.536, 331.6)}$ = 3.198 , P=0.0021; interaction $F_{(19, 836)}$ = 1.363, P=0.1369) and males (repeated measures two-way ANOVA for M SA Active NP vs. M SA Inactive NP over 20 sessions; NP $F_{(1, 38)}$ = 29.03, P<0.0001; session $F_{(2.536, 96.36)}$ = 2.246, P=0.0981; interaction $F_{(19, 722)}$ = 1.453, P=0.0954) without an effect between sexes (active NP: repeated measures two-way ANOVA for F SA Active NP vs. M SA Active NP over 20 sessions; NP $F_{(1, 41)}$ = 0.2039, P=0.6540; session $F_{(4.347, 178.2)}$ = 2.754, P=0.0258; interaction $F_{(19, 779)}$ = 1.377, P=0.1301 – inactive NP: repeated measures two-way ANOVA for F SA Inactive NP vs. M SA Inactive NP over 20 sessions; NP $F_{(1, 41)}$ = 3.758, P=0.0595; session $F_{(5.805, 238.0)}$ = 2.710, P=0.0595; interaction $F_{(19, 779)}$ = 1.517, P=0.0723).

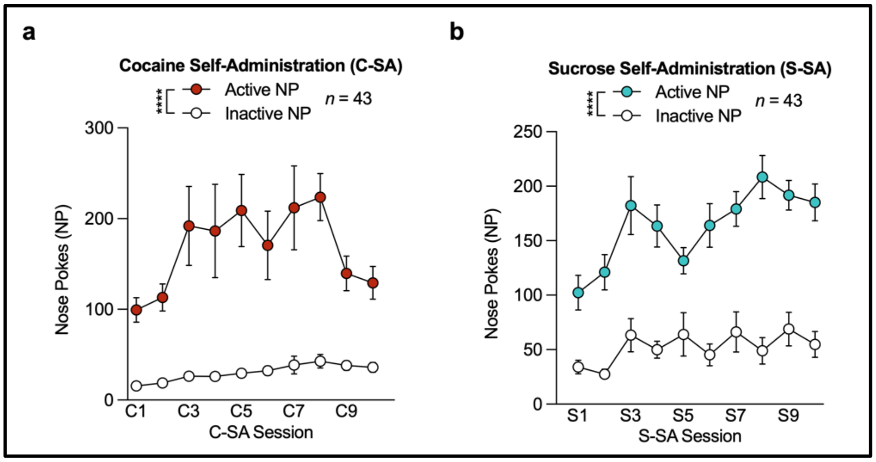


**Supplemental Figure 2 | Reward intake graphs. a,b.** Cocaine (**a,** C-SA) and sucrose (**b**, S-SA) self-administration sessions separated by cocaine (repeated measures two-way ANOVA for C-SA Active NP vs. C-SA Inactive NP over 20 sessions; NP $F_{(1, 84)} = 37.00$, $P<0.0001$; session $F_{(2.079, 174.6)} = 3.373$, $P=0.0348$; interaction $F_{(9, 756)} = 2.141$, $P=0.0243$) or sucrose (repeated measures two-way ANOVA for S-SA Active NP vs. S-SA Inactive NP over 20 sessions; NP $F_{(1, 84)} = 78.17$, $P<0.0001$; session $F_{(4.613, 387.5)} = 5.112$, $P=0.0002$; interaction $F_{(9, 756)} = 2.068$ , $P=0.0301$).

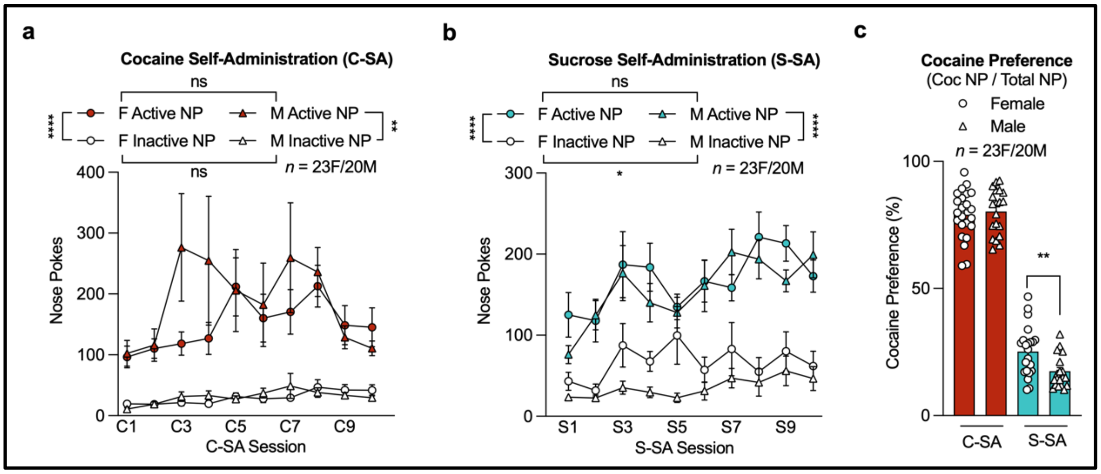


**Supplemental Figure 3 | Sex-specific self-administration and cocaine preference graphs. a.** Sex-specific analysis for C-SA (female: repeated measures two-way ANOVA for F C-SA Active NP vs. F C-SA Inactive NP over 20 sessions; NP $F_{(1, 44)}$ = 50.16, P<0.0001; session $F_{(4.496, 197.8)}$ = 2.689, P=0.0269; interaction $F_{(9, 396)}$ = 1.333, P=0.2177 – male: repeated measures two-way ANOVA for M C-SA Active NP vs. M C-SA Inactive NP over 20 sessions; NP $F_{(1, 38)}$ = 12.29, P=0.0012; session $F_{(1.511, 57.41)}$ = 2.513, P=0.1305; interaction $F_{(9, 342)}$ = 1.682, P=0.0919 – female/male active NP: repeated measures two-way ANOVA for F C-SA Active NP vs. M C-SA Active NP over 20 sessions; NP $F_{(1, 41)}$ = 0.6792, P=0.4146; session $F_{(2.024, 82.97)}$ = 2.907 , P=0.0596; interaction $F_{(9, 369)}$ = 1.490, P=0.1496 – female/male inactive NP: repeated measures two-way ANOVA for F C-SA Inactive NP vs. M C-SA Inactive NP over 20 sessions; NP $F_{(1, 41)}$ = 0.03314, P=0.8564; session $F_{(4.257, 174.5)}$ = 3.108, P=0.0148; interaction $F_{(9, 369)}$ = 1.236, P=0.2716). **b.** Sex-specific analysis for sucrose S-SA (female: repeated measures two-way ANOVA for F S-SA Active NP vs. F S-SA Inactive NP over 20 sessions; NP $F_{(1, 44)}$ = 26.13, P<0.0001; session $F_{(4.082, 179.6)}$ = 2.383, P=0.0520; interaction $F_{(9, 396)}$ = 1.425, P=0.1752 – male: repeated measures two-way ANOVA for M S-SA Active NP vs. M S-SA Inactive NP over 20 sessions; NP $F_{(1, 38)}$ = 77.63, P<0.0001; session $F_{(4.696, 178.5)}$ = 4.224, P=0.0015; interaction $F_{(9, 342)}$ = 1.739, P=0.0790 – female/male active NP: repeated measures two-way ANOVA for F S-SA Active NP vs. M S-SA Active NP over 20 sessions; NP $F_{(1, 41)}$ = 0.3130, P=0.5789; session $F_{(4.001, 164.0)}$ = 4.628, P=0.0014; interaction $F_{(9, 369)}$ = 0.9657, P=0.4681 – female/male inactive NP repeated measures two-way ANOVA for F S-SA Inactive NP vs. M S-SA Inactive NP over 20 sessions; NP $F_{(1, 41)}$ = 4.511, P=0.0398; session $F_{(3.873, 158.8)}$ = 1.446, P=0.2226; interaction $F_{(9, 369)}$ = 0.8511, P=0.5695). **c.** Sex-specific analysis of cocaine preference (cocaine NP / sucrose NP * 100) during C-SA and S-SA (unpaired t-test for F S-SA Cocaine Preference vs. M S-SA Cocaine Preference; P=0.0040).

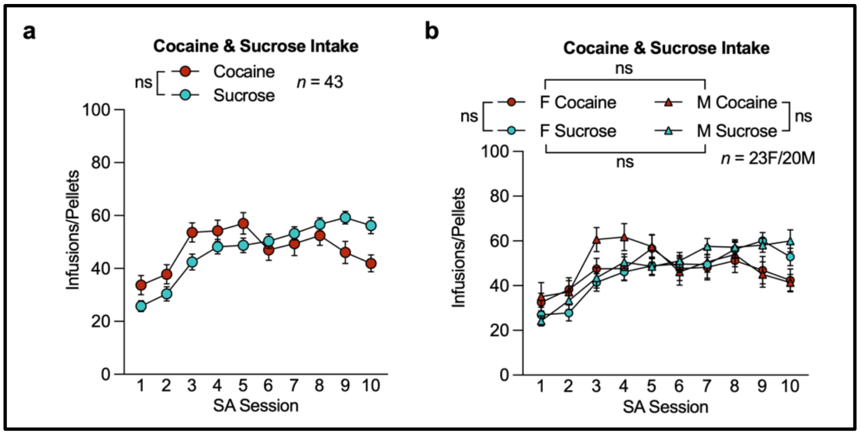


**Supplemental Figure 4 | Sex-specific reward intake graphs. a.** Reward-intake of cocaine (0.5 mg/kg/infusion) injections or sucrose (15 mg) pellets (max = 100 injections/pellets) upon association with active reward-specific nose poke (NP) ports showed no differentiation between (repeated measures two-way ANOVA for Cocaine Infusions vs. Sucrose Pellets over 20 sessions; NP $F_{(1, 84)}$ = 0.005, P=0.9446; session $F_{(4.805, 403.6)}$ = 17.59, P<0.0001; interaction $F_{(9, 756)}$ = 5.347, P<0.0001). **b.** Sex-specific analysis showed no differences between intake of cocaine and sucrose for females (repeated measures two-way ANOVA for F Cocaine Infusions vs. F Sucrose Pellets over 20 sessions; NP $F_{(1, 44)}$ = 0.00045, P=0.9832; session $F_{(4.269, 187.8)}$ = 9.184, P<0.0001; interaction $F_{(9, 396)}$ = 2.065, P=0.0316) and males (repeated measures two-way ANOVA for M Cocaine Infusions vs. M Sucrose Pellets over 20 sessions; NP $F_{(1, 38)}$ = 0.01675, P=0.8977; session $F_{(4.638, 176.2)}$ = 9.083 , P<0.0001; interaction $F_{(9, 342)}$ = 3.911, P<0.0001). Comparing intake of cocaine between sexes showed no difference (repeated measures two-way ANOVA for F Cocaine Infusions vs. M Cocaine Infusions over 20 sessions; NP $F_{(1, 41)}$ = 0.4724, P=0.4957; session $F_{(4.947, 202.8)}$ = 5.182, P=0.0002; interaction $F_{(9, 369)}$ = 0.7462, P=0.6664). Comparing intake of sucrose between sexes showed no difference (repeated measures two-way ANOVA for F Sucrose Pellets vs. M Sucrose Pellets over 20 sessions; NP $F_{(1, 41)}$ = 0.5292, P=0.4711; session $F_{(3.598, 147.5)}$ = 26.71, P<0.0001; interaction $F_{(9, 369)}$ = 0.7441, P=0.6684).

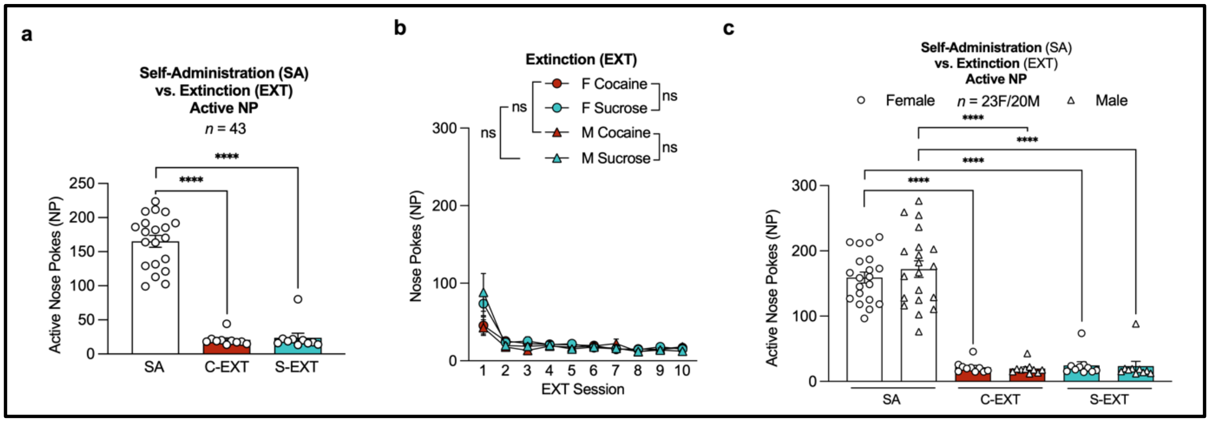


**Supplemental Figure 5 | Sex-specific extinction graphs. a.** During EXT of cocaine (C-EXT) and sucrose (S-EXT), mice showed a decrease in active NP interaction when compared to cocaine (mixed-effect repeated measures two-way ANOVA for SA Active NP vs. C-EXT Active NP over 20 sessions; NP $F_{(1, 84)}$ = 64.52, P<0.0001; session $F_{(3.311, 206.5)}$ = 2.615, P=0.0466) or sucrose SA (S-SA) sessions (mixed-effect repeated measures two-way ANOVA for SA Active NP vs. S-EXT Active NP over 20 sessions; NP $F_{(1, 84)}$ = 60.81, P<0.0001; session $F_{(3.389, 211.4)}$ = 2.495, P=0.0536). **b.** Sex-specific plot for extinction show no differentiation between cocaine and sucrose active NP for females (repeated measures two-way ANOVA for F EXT Cocaine NP vs. F EXT Sucrose NP over 20 sessions; NP $F_{(1, 44)}$ = 0.4072, P=0.5267; session $F_{(1.782, 78.39)}$ = 17.28, P<0.0001; interaction $F_{(9, 396)}$ = 2.001, P=0.0380) and males (repeated measures two-way ANOVA for M EXT Cocaine NP vs. M EXT Sucrose NP over 20 sessions; NP $F_{(1, 38)}$ = 0.5938, P=0.4457; session $F_{(1.624, 61.73)}$ = 12.61, P<0.0001; interaction $F_{(9, 342)}$ = 2.916, P=0.0024) without an effect between sexes for cocaine NP (repeated measures two-way ANOVA for F EXT Cocaine NP vs. M EXT Cocaine NP over 20 sessions; NP $F_{(1, 41)}$ = 0.2101, P=0.6491; session $F_{(2.332, 95.61)}$ = 9.430, P<0.0001; interaction $F_{(9, 369)}$ = 0.6856, P=0.7220) and sucrose NP (repeated measures two-way ANOVA for F EXT Sucrose NP vs. M EXT Sucrose NP over 20 sessions; NP $F_{(1, 41)}$ = 0.05162, P=0.8214; session $F_{(1.473, 60.38)}$ = 19.73, P<0.0001; interaction $F_{(9, 369)}$ = 0.4548, P=0.9041). **c.** When comparing active NP interactions during SA to C-EXT, there was a significant decrease in the active NP during C-EXT for females (mixed-effect repeated measures two-way ANOVA for F SA Active NP vs. F C-EXT Active NP over 20 sessions; NP $F_{(1, 44)}$ = 48.42 , P<0.0001; session $F_{(5.194, 170.9)}$ = 2.164, P=0.0578) and males (mixed-effect repeated measures two-way ANOVA for M SA Active NP vs. M C-EXT Active NP over 20 sessions; NP $F_{(1, 38)}$ = 23.88, P<0.0001; session $F_{(1.649, 46.96)}$ = 1.809, P=0.1803). Comparing active NP interactions during SA to S-EXT, there was a significant decrease in the active NP during S-EXT for females (mixed-effect repeated measures two-way ANOVA for F SA Active NP vs. F S-EXT Active NP over 20 sessions; NP $F_{(1, 44)}$ = 45.97, P<0.0001; session $F_{(5.291, 174.1)}$ = 2.099, P=0.0639) and males (mixed-effect repeated measures two-way ANOVA for M SA Active NP vs. M S-EXT Active NP over 20 sessions; NP $F_{(1, 38)}$ = 22.39, P<0.0001; session $F_{(1.698, 48.34)}$ = 1.696, P=0.1975).

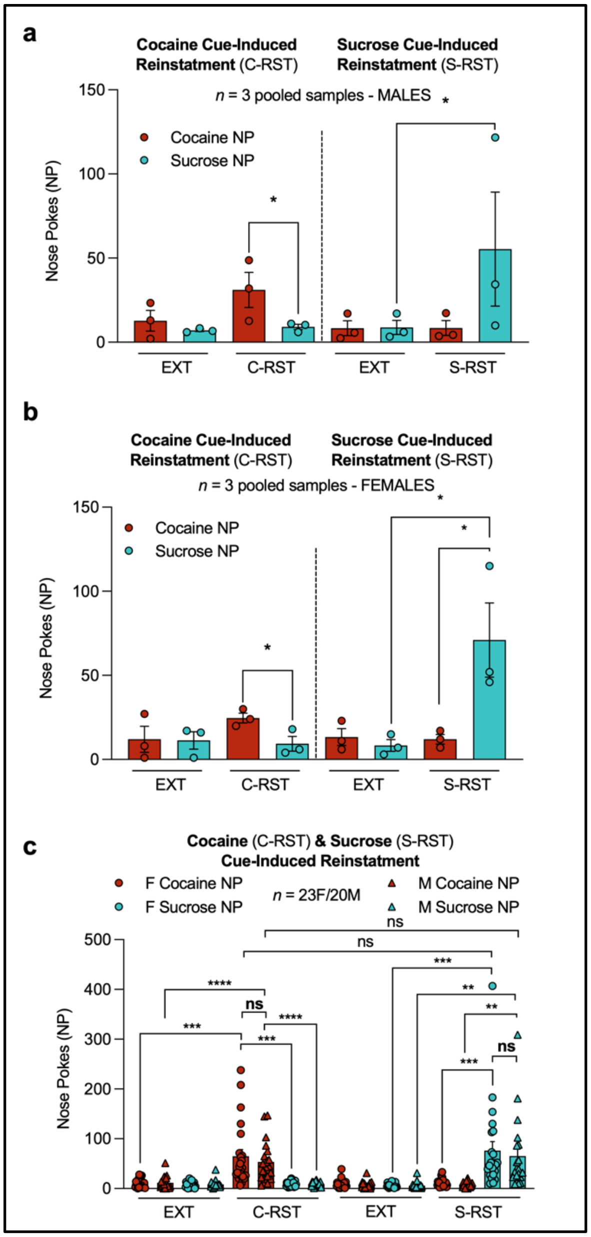


**Supplemental Figure 6 | Sex-specific reinstatements graphs. a.** Plot of averaged pooled male sample reinstatement grouped by RNA pooling schematic of 3 mice per sample per three replicates. Male pooled samples displayed significant differentiation between the cocaine NP and sucrose NP during cocaine reinstatement (C-RST) (paired t-test for Average of Male Sample Pools: C-RST Cocaine NP vs. C-RST Sucrose NP; P=0.0344). Male pooled samples displayed significant differentiation of sucrose NP interaction between extinction (EXT) and sucrose reinstatement (S-RST) (paired t-test for Average of Male Sample Pools: S-RST Sucrose NP vs. S-RST Cocaine NP; P=0.0592). **b.** Plot of averaged pooled female sample reinstatement grouped by RNA pooling schematic of 3 mice per sample per three replicates. Female pooled samples displayed significant differentiation between the cocaine NP and sucrose NP during C-RST (paired t-test for Average of Female Sample Pools: C-RST Cocaine NP vs. C-RST Sucrose NP; P=0.0237) and S-RST (paired t-test for Average of Female Sample Pools: S-RST Sucrose NP vs. S-RST Cocaine NP; P=0.0136). Female pooled samples displayed significant differentiation of sucrose NP interaction between EXT and S-RST (paired t-test for Average of Female Sample Pools: S-RST Sucrose NP vs. S-EXT Sucrose NP; P=0.047). **c.** Sex-specific plot for C-RST shows higher interaction with the cocaine NP during C-RST compared to the sucrose NP for females (paired t-test for F C-RST Cocaine NP vs. F C-RST Sucrose NP; P=0.0002) and males (paired t-test for M C-RST Cocaine NP vs. M C-RST Sucrose NP; P<0.0001). There was a significant increase in interactions with the cocaine NP during C-RST compared to the last day of EXT for females (paired t-test for F C-RST Cocaine NP vs. F C-EXT Cocaine NP; P=0.0009) and males (paired t-test for M C-RST Cocaine NP vs. M C-EXT Cocaine NP; P=0.0018). Sex-specific plot for S-RST shows higher interaction with the sucrose NP during S-RST compared to the cocaine NP for females (paired t-test for F S-RST Sucrose NP vs. F S-RST Cocaine NP; P=0.0005) and males (paired t-test for M S-RST Sucrose NP vs. M S-RST Cocaine NP; P<0.0001). There was a significant increase in interactions with the sucrose NP during S-RST compared to the last day of extinction for females (paired t-test for F S-RST Sucrose NP vs. F S-EXT Sucrose NP; P=0.0006) and males (paired t-test for M S-RST Sucrose NP vs. M S-EXT Sucrose NP; P=0.0011). Comparing active NP interactions in C-RST and S-RST showed no difference for females (paired t-test for F C-RST Cocaine NP vs. F S-RST Sucrose NP; P=0.3923) and males (paired t-test for M C-RST Cocaine NP vs. M S-RST Sucrose NP; P=0.4379). Comparing the interaction of active NP during C-RST between males and females showed no difference (unpaired t-test for F C-RST Cocaine NP vs. M C-RST Cocaine NP; P=0.4866). Comparing the interaction of active NP during S-RST between males and females showed no difference (unpaired t-test for F S-RST Sucrose NP vs. M S-RST Sucrose NP; P=0.6535).

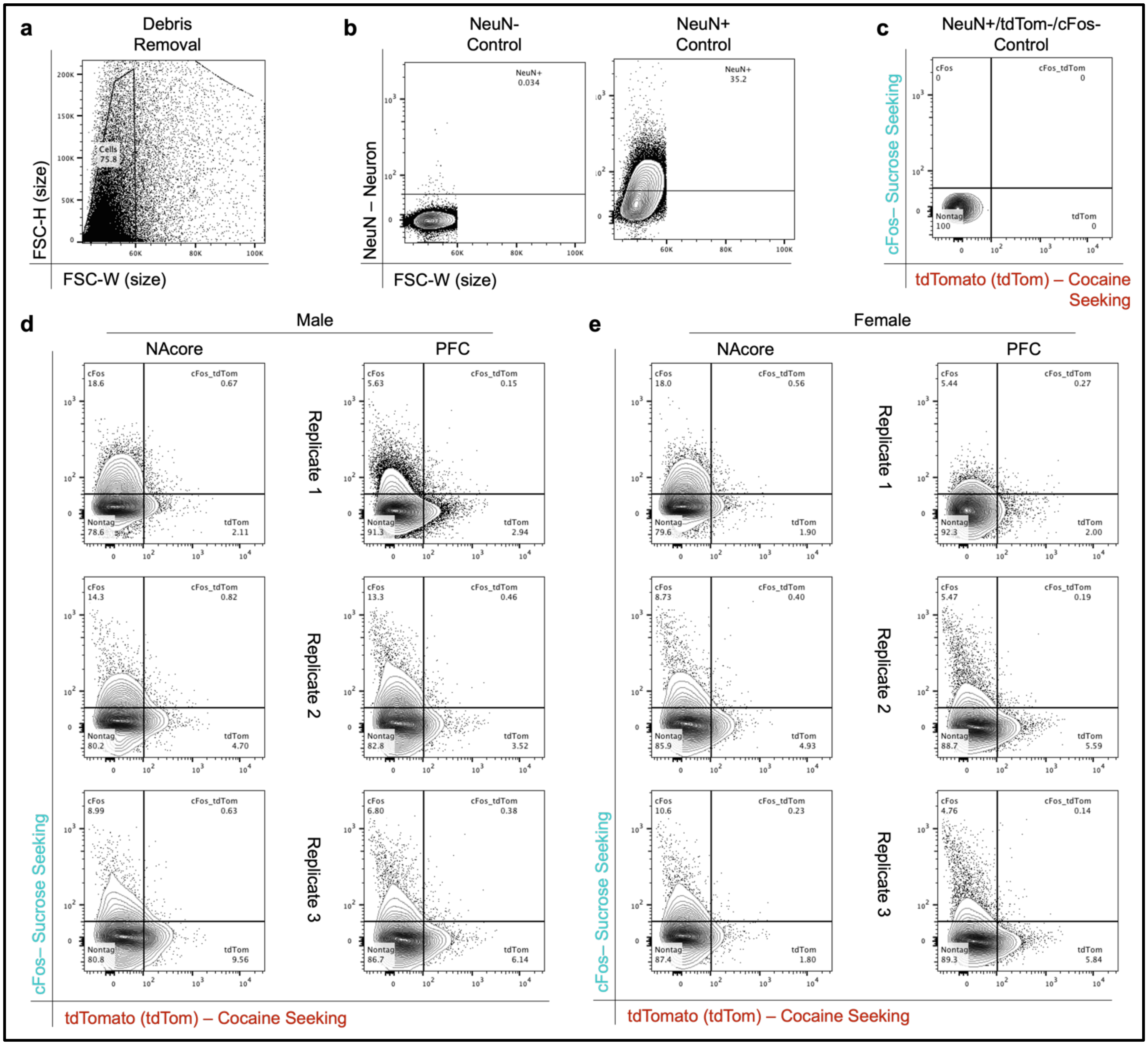


**Supplemental Figure 7 | Fluorescent-activated cell sorting (FACS) gating strategies and flow cytometric analysis of sample replicates. a.** Proof of concept for FACS debris removal from a sample using the size parameters forward scatter width (FSC-W; x-axis) and forward scatter height (FSC-H; y-axis) to filter out unwanted larger and smaller events. **b.** We further gated cells positive for a neuronal marker (NeuN+) in comparison to our negative control (NeuN-) to gate for neurons to use in downstream ensemble-specific quadrant plots. **c.** We used transgenic mice who went through behavioral conditioning paradigm but lacked the induction of tagging scheme as a control for quadrant plot gating strategy from gated NeuN+ cells in b to gate for cocaine-seeking (tdTom; x-axis) and sucrose-seeking (cFos; y-axis) markers. **d.** Male flow cytometric quadrant plots subcategorized into the nucleus accumbens core (NAcore) and prefrontal cortex (PFC) regions for FACS-sorted ensemble specific samples. **e.** Female flow cytometric quadrant plots subcategorized into the nucleus accumbens core (NAcore) and prefrontal cortex (PFC) regions for FACS-sorted ensemble specific samples. For males and females, quadrant analysis allowed separation of cocaine-seeking neurons (bottom-right; tdTom+/cFos-), sucrose-seeking neurons (top left; cFos+/tdTom-) and overlapping neurons between cocaine-seeking and sucrose-seeking (top right: tdTom+/cFos+) in comparison to non-ensemble (bottom-left; tdTom-/cFos-) neurons. Quadrant plots are shown for replicates 1-3 with replicate 3 used as representation in Figure 1f,g. f-i. Comparisons for quantification of cocaine-seeking (C), sucrose-seeking (S), and overlapping (O) ensemble sizes (% of tagged ensemble-specific neurons per 1,000,000 events).

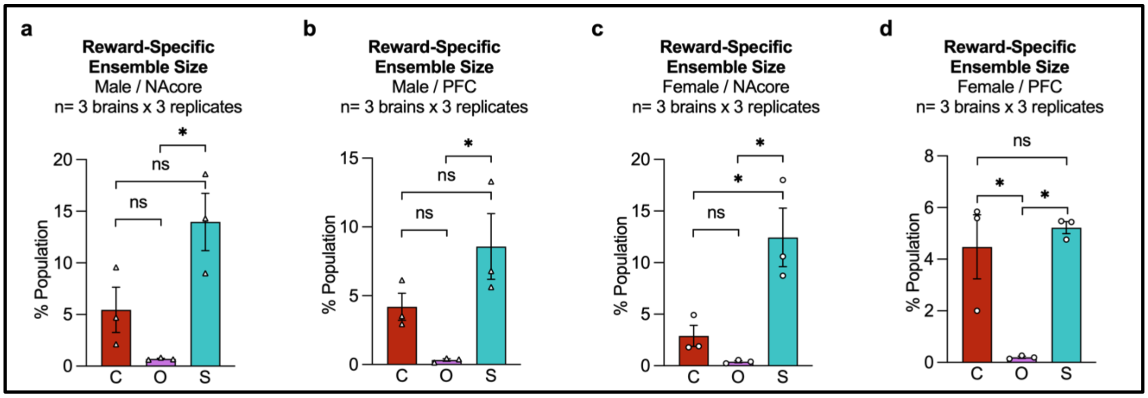


**Supplemental Figure 8 | Flow cytometric analysis of ensemble sizes. a.** Within the male NAcore, there was an overall significant difference in ensemble sizes and replicates (repeated measures two-way ANOVA for M Cocaine NAcore vs. M Sucrose NAcore vs. M Overlap NAcore: Reward vs. Replicate; reward $F_{(2, 4)}$ = 7.303, P=0.0462; replicate $F_{(2, 4)}$ = 0.02303, P=0.04462), a significant difference between the size of sucrose-seeking and overlapping ensemble (multiple comparisons: repeated measures two-way ANOVA for M Sucrose NAcore vs. M Overlap NAcore; P=0.0419), no significant differences between the size of cocaine-seeking and sucrose-seeking ensembles (multiple comparisons: repeated measures two-way ANOVA for M Cocaine NAcore vs. M Sucrose NAcore; P=0.11475), and no significant difference between the size of the cocaine-seeking ensemble and overlapping ensemble (multiple comparisons: repeated measures two-way ANOVA for M Cocaine NAcore vs. M Overlap NAcore; P=0.4435). **b.** Within the female NAcore, there was an overall significant difference in ensemble sizes (repeated measures two-way ANOVA for F Cocaine NAcore vs. F Sucrose NAcore vs. F Overlap NAcore: Reward vs. Replicate; reward $F_{(2,4)}$ = 7.354, P=0.0457; replicate $F_{(2, 4)}$ = 0.8809, P=0.4820), a significant difference between the size of sucrose-seeking and overlapping ensemble (multiple comparisons: repeated measures two-way ANOVA for F Sucrose NAcore vs. F Overlap NAcore; P=0.0398), no significant differences between the size of cocaine-seeking and sucrose-seeking ensembles (multiple comparisons: repeated measures two-way ANOVA for F Cocaine NAcore vs. F Sucrose NAcore; P=0.2196), and no significant difference between the size of the cocaine-seeking ensemble and overlapping ensemble (multiple comparisons: repeated measures two-way ANOVA for F Cocaine NAcore vs. F Overlap NAcore; P=0.2808). **c.** Within the male PFC, there was an overall significant difference in ensemble sizes (repeated measures two-way ANOVA for M Cocaine PFC vs. M Sucrose PFC vs. M Overlap PFC: Reward vs. Replicate; reward $F_{(2, 4)}$ =11.33, P=0.0225; replicate $F_{(2, 4)}$ = 0.5409, P=0.6196), a significant difference between the size of cocaine-seeking and sucrose-seeking ensembles (multiple comparisons: repeated measures two-way ANOVA for M Cocaine PFC vs. M Sucrose PFC; P=0.0493), a significant difference between the size of sucrose-seeking and overlapping ensemble (multiple comparisons: repeated measures two-way ANOVA for M Sucrose PFC vs. M Overlap PFC; P=0.0233), and no significant differences between the size of cocaine-seeking and overlapping ensembles (multiple comparisons: repeated measures two-way ANOVA for M Cocaine PFC vs. M Overlap PFC; P=0.6537). **d.** Within the female PFC, there was an overall significant difference in ensemble sizes (repeated measures two-way ANOVA for F Cocaine PFC vs. F Sucrose PFC vs. F Overlap PFC: Reward vs. Replicate; reward $F_{(2,4)}$ = 12.38, P=0.0193; replicate $F_{(2,4)}$ = 0.6856, P=0.5546), a significant difference between the size of cocaine-seeking and overlapping ensembles (multiple comparisons: repeated measures two-way ANOVA for F Cocaine PFC vs. F Overlap PFC; P=0.0368), a significant difference between the size of sucrose-seeking and overlapping ensemble (multiple comparisons: repeated measures two-way ANOVA for F Sucrose PFC vs. F Overlap PFC; P=0.0216), and no significant differences between the size of cocaine-seeking and overlapping ensembles (multiple comparisons: repeated measures two-way ANOVA for F Cocaine PFC vs. F Sucrose PFC; P=0.7841).

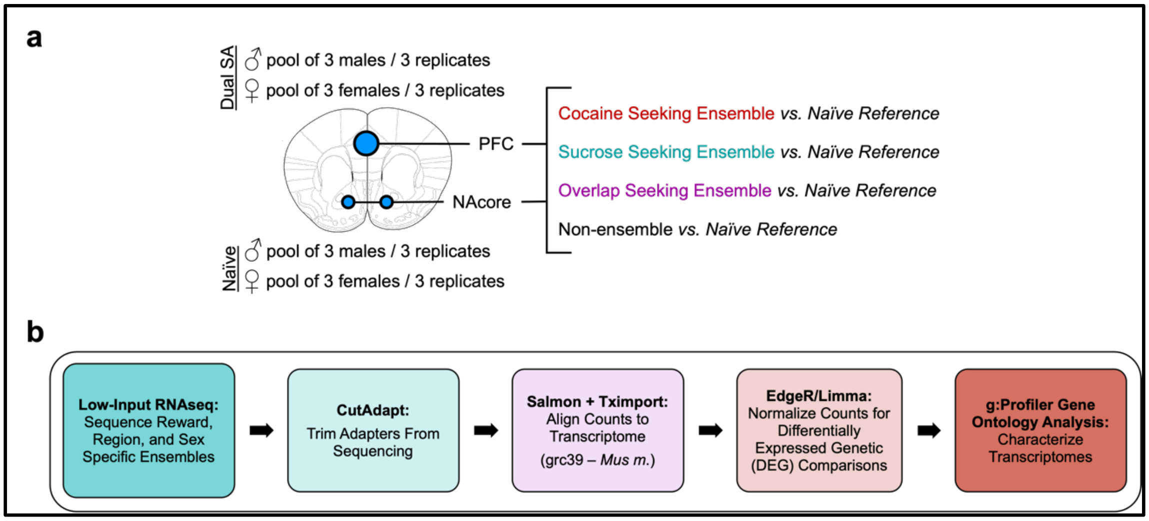


**Supplemental Figure 9 | Overview of transcriptomic comparison analysis. a,b.** Bioinformatic workflow used to determine differentially expressed genes (DEG) for each FACS-sorted reward-specific seeking ensemble in the NAcore and PFC in comparison to naïve females and males that did not undergo dual cocaine and sucrose self-administration (SA).

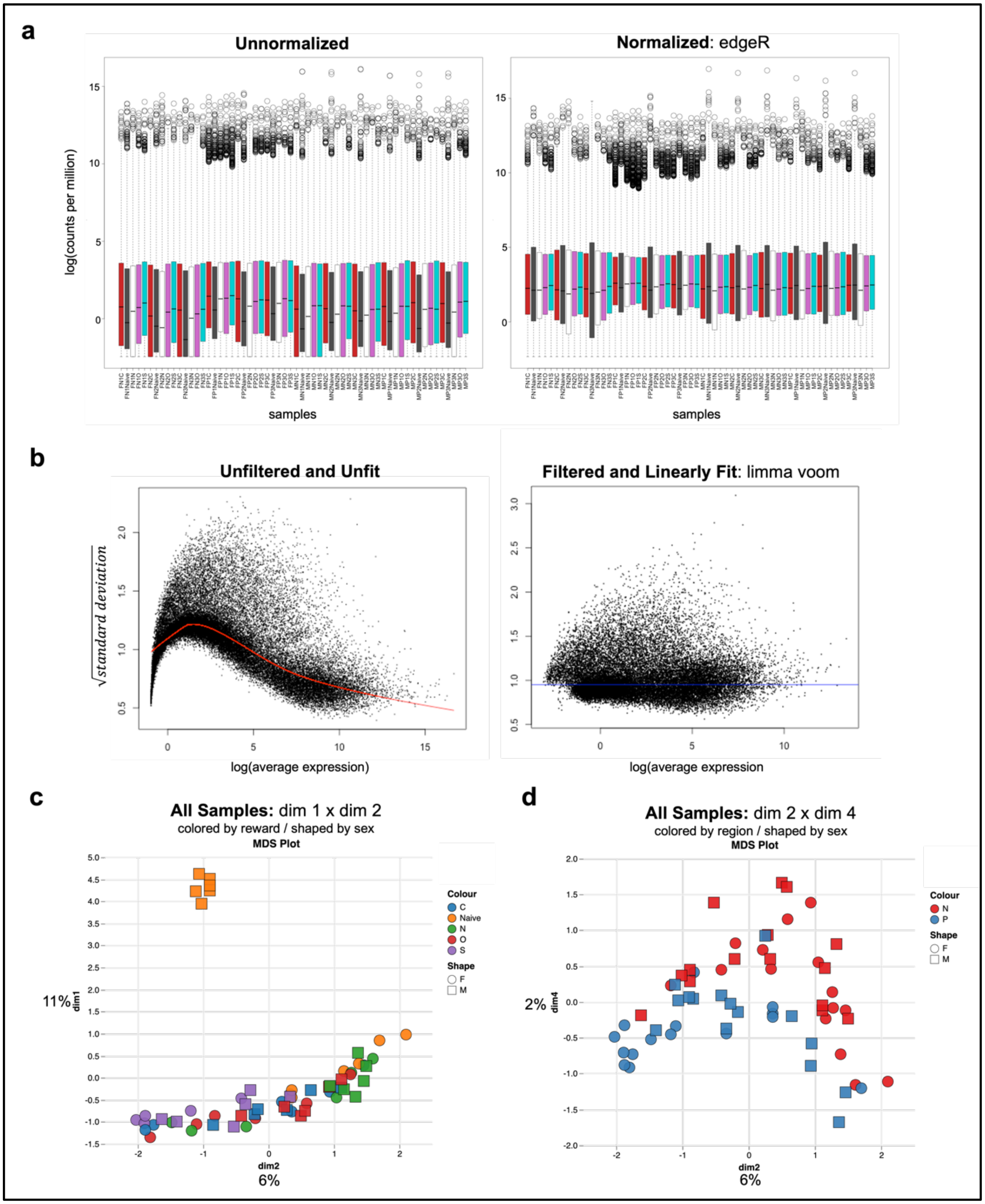


**Supplemental Figure 10 | Plots for quality control of count data for all samples. a.** Unnormalized (left panel) and edgeR normalized (right panel) boxplots scaled by log(counts per million) (y-axis) and colored by sample type (cocaine-seeking ensemble = red; sucrose-seeking ensemble = aqua; overlapping ensemble = purple; non-ensemble = grey; naïve = white) (x-axis) for all samples. **b.** Scatter plots of log(average expression of replicates) (x-axis) and the square root of standard deviation (y-axis) comparing unfiltered and unfit (left panel) counts and limma-based filtered and linearly fit counts (right panel) for all samples. **c.** Multidimensional scaling plots for all samples colored by reward ensemble (cocaine-seeking ensemble = C; sucrose-seeking ensemble= S; overlapping ensemble= O; non-ensemble = N) and shaped by sex in the nucleus accumbens core and prefrontal cortex. Percent of variation is included on each axis representative of dim1 or dim2. **d.** Multidimensional scaling plots for all samples colored by region and shaped by sex (male = M; female = F) in the nucleus accumbens core (N) and prefrontal cortex (P). Percent of variation is included on each axis representative of dim4 or dim2.

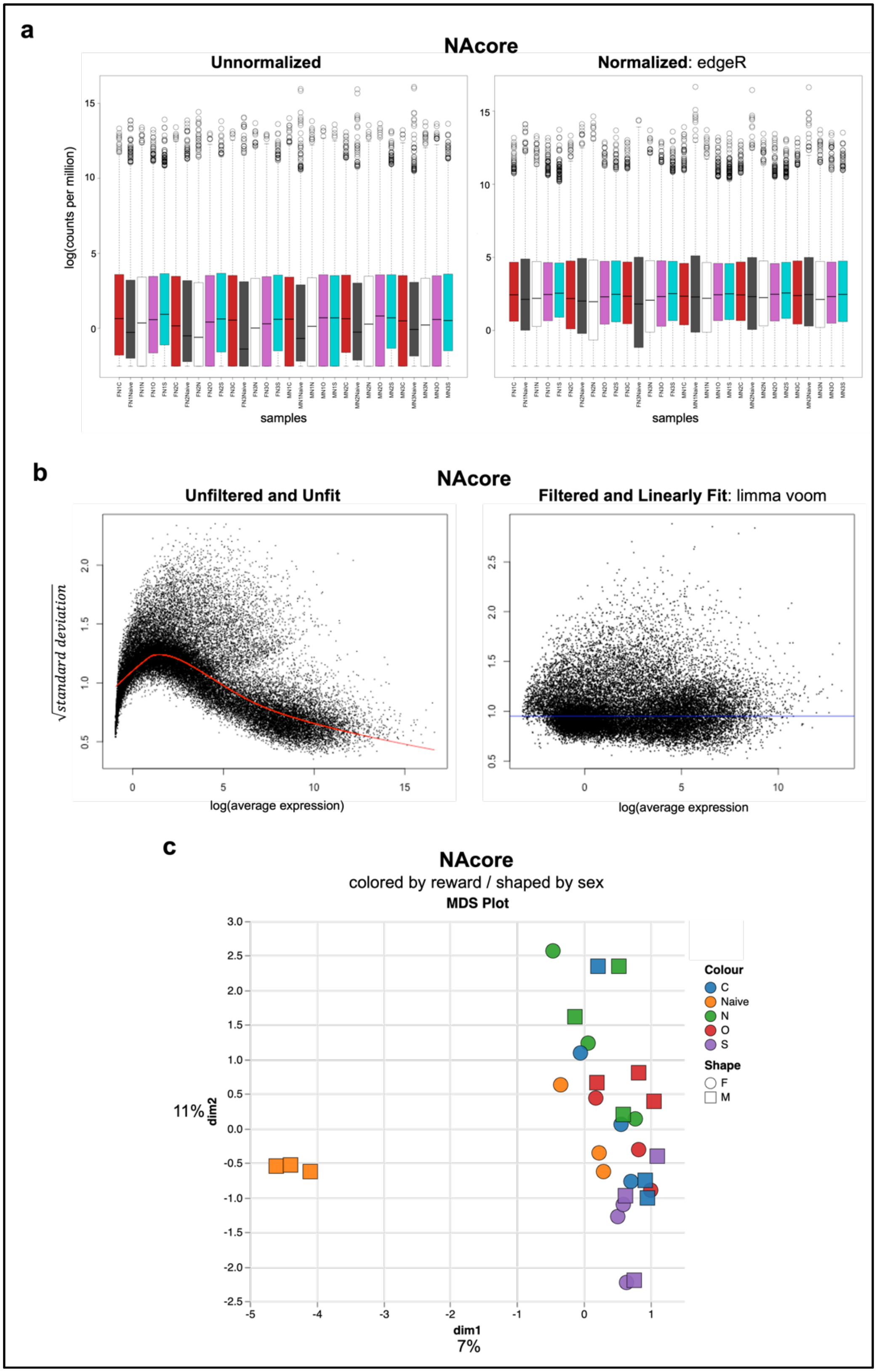


**Supplemental Figure 11 | Plots for quality control of count data for samples in the NAcore. a.** Unnormalized (left panel) and edgeR normalized (right panel) boxplots scaled by log(counts per million) (y-axis) and colored by sample type (cocaine-seeking ensemble = red; sucrose-seeking ensemble = aqua; overlapping ensemble = purple; non-ensemble = grey; naïve = white) (x-axis) for samples in the NAcore. **b.** Scatter plots of log(average expression of replicates) (x-axis) and the square root of standard deviation (y-axis) comparing unfiltered and unfit (left panel) counts and limma-based filtered and linearly fit counts (right panel) for samples in the NAcore. **c.** Multidimensional scaling plots for samples in the NAcore colored by reward and shaped by sex. Percent of variation is included on each axis representative of dim1 or dim2.

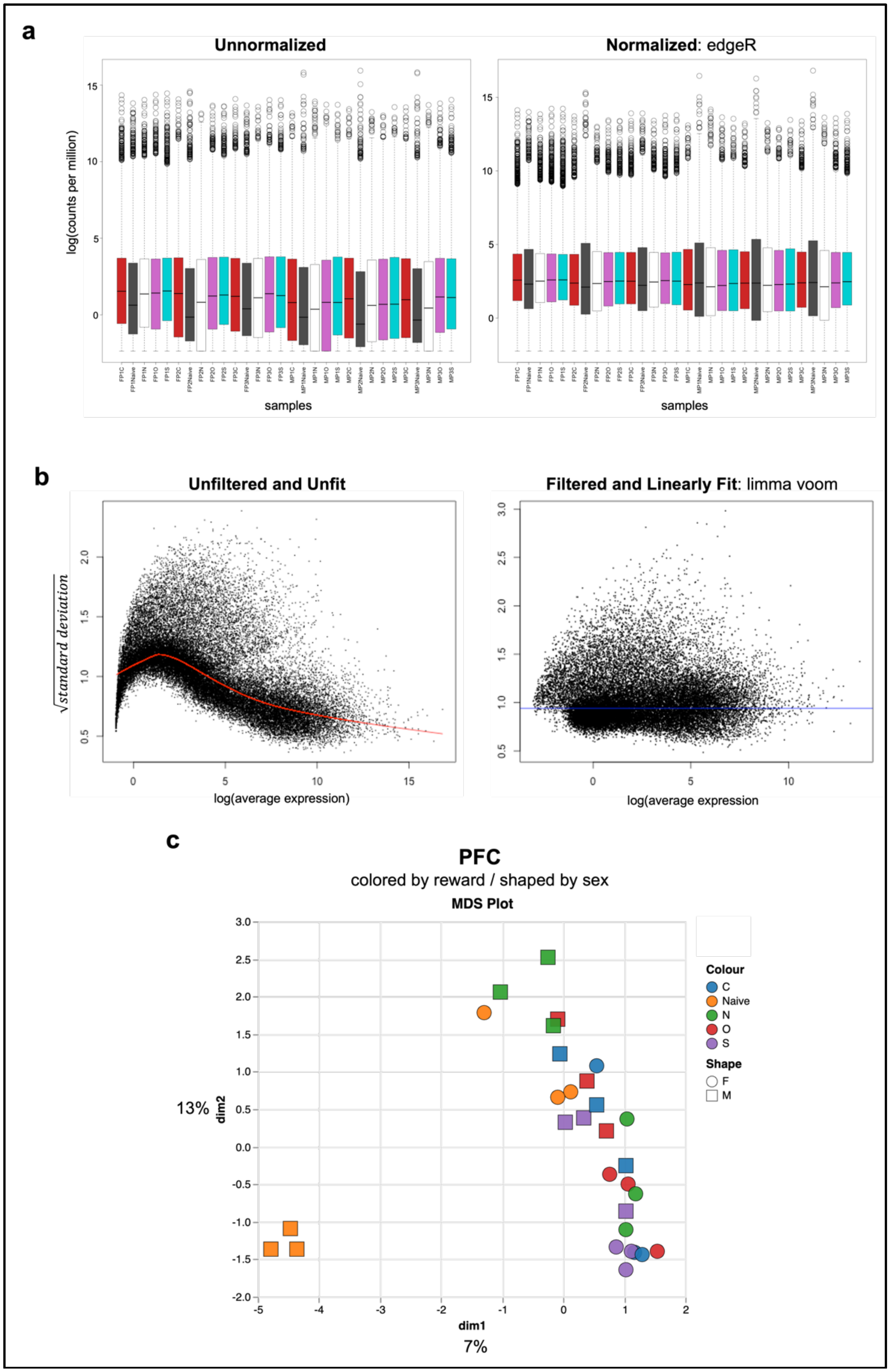


**Supplemental Figure 12 | Plots for quality control of count data for samples in the PFC. a.** Unnormalized (left panel) and edgeR normalized (right panel) boxplots scaled by log(counts per million) (y-axis) and colored by sample type (cocaine-seeking ensemble = red; sucrose-seeking ensemble = aqua; overlapping ensemble = purple; non-ensemble = grey; naïve = white) (x-axis) for samples in the PFC. **b.** Scatter plots of log(average expression of replicates) (x-axis) and the square root of standard deviation (y-axis) comparing unfiltered and unfit (left panel) counts and limma-based filtered and linearly fit counts (right panel) for samples in the PFC. **c.** Multidimensional scaling plots for samples in the PFC colored by reward and shaped by sex. Percent of variation is included on each axis representative of dim1 or dim2.
